# mRNA Vaccination Induces Durable Immune Memory to SARS-CoV-2 with Continued Evolution to Variants of Concern

**DOI:** 10.1101/2021.08.23.457229

**Authors:** Rishi R. Goel, Mark M. Painter, Sokratis A. Apostolidis, Divij Mathew, Wenzhao Meng, Aaron M. Rosenfeld, Kendall A. Lundgreen, Arnold Reynaldi, David S. Khoury, Ajinkya Pattekar, Sigrid Gouma, Leticia Kuri-Cervantes, Philip Hicks, Sarah Dysinger, Amanda Hicks, Harsh Sharma, Sarah Herring, Scott Korte, Amy E. Baxter, Derek A. Oldridge, Josephine R. Giles, Madison E. Weirick, Christopher M. McAllister, Moses Awofolaju, Nicole Tanenbaum, Elizabeth M. Drapeau, Jeanette Dougherty, Sherea Long, Kurt D’Andrea, Jacob T. Hamilton, Maura McLaughlin, Justine C. Williams, Sharon Adamski, Oliva Kuthuru, The UPenn COVID Processing Unit, Ian Frank, Michael R. Betts, Laura A. Vella, Alba Grifoni, Daniela Weiskopf, Alessandro Sette, Scott E. Hensley, Miles P. Davenport, Paul Bates, Eline T. Luning Prak, Allison R. Greenplate, E. John Wherry

## Abstract

SARS-CoV-2 mRNA vaccines have shown remarkable efficacy, especially in preventing severe illness and hospitalization. However, the emergence of several variants of concern and reports of declining antibody levels have raised uncertainty about the durability of immune memory following vaccination. In this study, we longitudinally profiled both antibody and cellular immune responses in SARS-CoV-2 naïve and recovered individuals from pre-vaccine baseline to 6 months post-mRNA vaccination. Antibody and neutralizing titers decayed from peak levels but remained detectable in all subjects at 6 months post-vaccination. Functional memory B cell responses, including those specific for the receptor binding domain (RBD) of the Alpha (B.1.1.7), Beta (B.1.351), and Delta (B.1.617.2) variants, were also efficiently generated by mRNA vaccination and continued to increase in frequency between 3 and 6 months post-vaccination. Notably, most memory B cells induced by mRNA vaccines were capable of cross-binding variants of concern, and B cell receptor sequencing revealed significantly more hypermutation in these RBD variant-binding clones compared to clones that exclusively bound wild-type RBD. Moreover, the percent of variant cross-binding memory B cells was higher in vaccinees than individuals who recovered from mild COVID-19. mRNA vaccination also generated antigen-specific CD8+ T cells and durable memory CD4+ T cells in most individuals, with early CD4+ T cell responses correlating with humoral immunity at later timepoints. These findings demonstrate robust, multi-component humoral and cellular immune memory to SARS-CoV-2 and current variants of concern for at least 6 months after mRNA vaccination. Finally, we observed that boosting of pre-existing immunity with mRNA vaccination in SARS-CoV-2 recovered individuals primarily increased antibody responses in the short-term without significantly altering antibody decay rates or long-term B and T cell memory. Together, this study provides insights into the generation and evolution of vaccine-induced immunity to SARS-CoV-2, including variants of concern, and has implications for future booster strategies.

**GRAPHICAL ABSTRACT:** 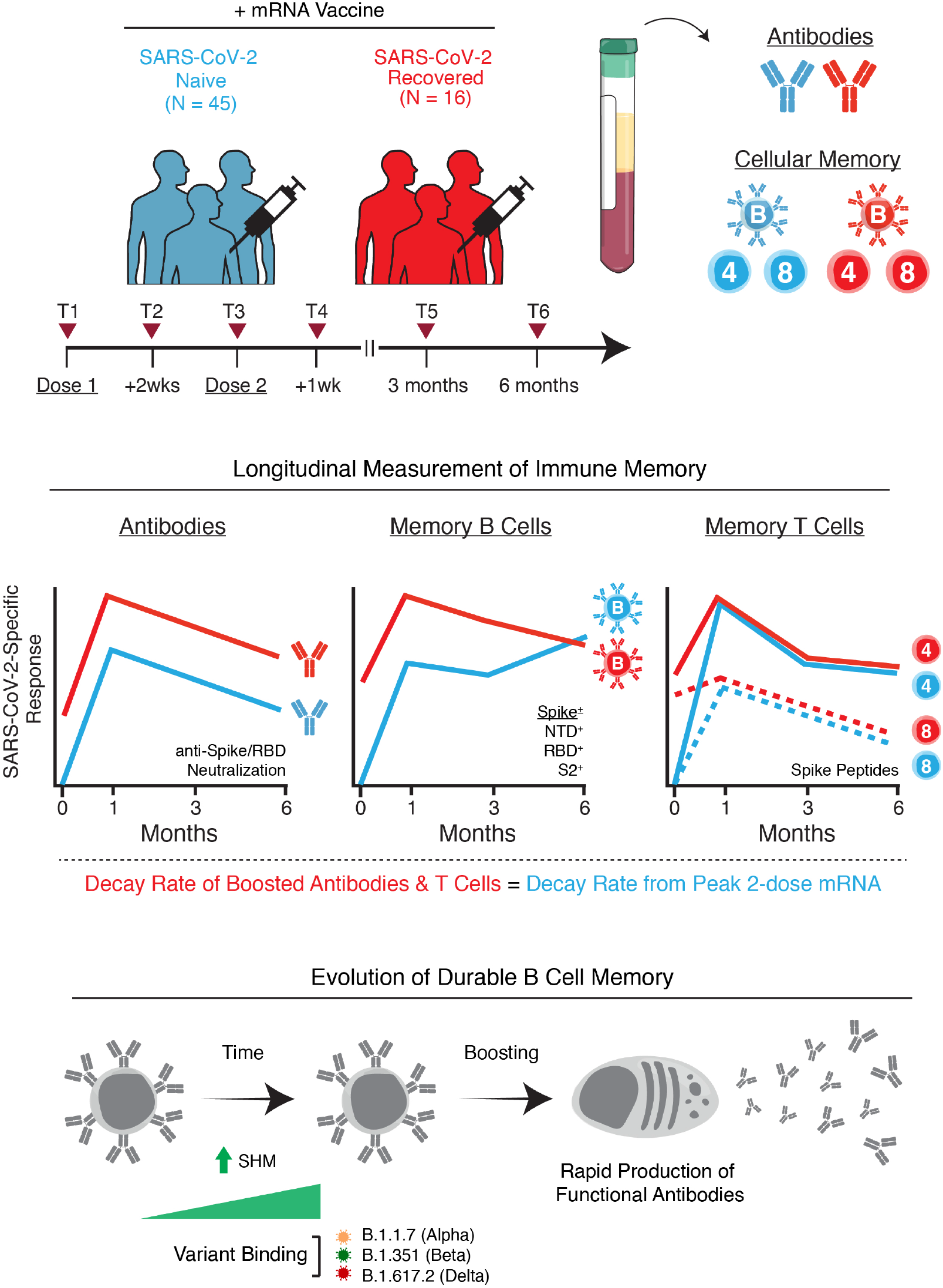

## INTRODUCTION

The coronavirus disease 2019 (COVID-19) pandemic continues to result in significant morbidity and mortality worldwide. Community-level immunity, acquired through natural infection or vaccination, is necessary to control the pandemic as the virus continues to circulate (*1*). mRNA vaccines encoding a stabilized version of the full-length SARS-CoV-2 Spike protein are being widely administered and clinical trial data demonstrate up to 95% efficacy in preventing symptomatic COVID-19 (*2, 3*). These vaccines induce potent humoral immune responses, with neutralizing antibody titers emerging as a correlate of protection (*4–6*). Current evidence suggests that circulating antibodies persist for at least 6 months post-vaccination (*7*), though there is some decay from peak levels achieved after the second dose. This decline from peak antibody levels may be associated with an increase in infections over time compared to the initial months post-vaccination (*8, 9*). However, other data indicate that vaccine-induced immunity remains highly effective at preventing severe disease, hospitalization, and death even at later timepoints when antibody levels may decline (*10*).

In addition to the production of antibodies, an effective immune response requires the generation of long-lived memory B and T cells. mRNA vaccines induce robust germinal center responses in humans (*11*), resulting in memory B cells that are specific for both the full-length SARS-CoV-2 Spike protein and the Spike receptor binding domain (RBD) (*12–14*). mRNA vaccination has also been shown to generate Spike-specific memory CD4+ and CD8+ T cell responses (*15–18*). Although antibodies are often correlates of vaccine efficacy, memory B cells and memory T cells are important components of the recall response to viral antigens and are a likely mechanism of protection, especially in the setting of infections in previously vaccinated individuals where antibodies alone do not provide sterilizing immunity (*19*). In such cases, memory B and T cells can be rapidly re-activated, resulting in enhanced control of initial viral replication and limiting viral dissemination in the host (*20, 21*). By responding and restricting viral infection within the first hours to days after exposure, cellular immunity can thereby reduce or even prevent symptoms of disease (i.e. preventing hospitalization and death) and potentially reduce the ability to spread virus to others (*22*).

Immunological studies of natural infection show that memory B and T cell responses appear to persist for at least 8 months post-symptom onset (*23, 24*). However, the durability of these populations of memory B and T cells following vaccination remains poorly understood. The emergence of several SARS-CoV-2 variants, including B.1.1.7 (Alpha), B.1.351 (Beta), and most recently B.1.617.2 (Delta), has also raised concerns about increased transmission and potential evasion from vaccine-induced immunity (*25–28*). As such, it is necessary to develop a more complete understanding of the trajectory and durability of immunological memory after mRNA vaccination, as well as how immune responses are affected by current variants of concern (VOCs). Moreover, the United States and other well-resourced countries have recently announced plans for a third vaccine booster dose, yet information on how pre-existing serological and cellular immunity to SARS-CoV-2 is boosted by mRNA vaccination remains limited. Specifically, it is unclear how different components of the immune response may benefit from boosting and whether boosting has any effect on the durability of these components. Exploring these questions will provide better insights into mRNA vaccine-induced immunological memory and will likely be relevant for future vaccine strategies, including recommendations for additional booster vaccine doses. Here, we examine these key questions by measuring SARS-CoV-2-specific antibody, memory B cell, and memory T cell responses through 6 months post-vaccination in a group of healthy subjects receiving 2 doses of mRNA vaccine compared with a group of SARS-CoV-2 recovered subjects where pre-existing immunity was boosted by mRNA vaccination.

## RESULTS AND DISCUSSION

### Cohort Design

We collected 342 longitudinal samples from 61 individuals receiving either the Pfizer BNT162b2 or Moderna mRNA-1273 SARS-CoV-2 vaccines at 6 timepoints (figure 1A), ranging from pre-vaccination baseline to 6 months post-vaccination. This study design allowed us to monitor the induction and maintenance of antigen-specific immune responses to the vaccine. Specifically, the inclusion of 1-, 3-, and 6-month samples enabled analysis of immune trajectories from peak responses after the second vaccine dose through establishment and maintenance of immunological memory. This cohort was divided into 2 groups based on evidence of a prior SARS-CoV-2 infection (N=45 SARS-CoV-2 naïve, N=16 SARS-CoV-2 recovered). Notably, the subjects with a prior infection allowed us to study the dynamics of boosting pre-existing immunity with mRNA vaccines. Demographic characteristics (age and sex) were balanced in both groups. Paired serum and peripheral blood mononuclear cell (PBMC) samples were collected on all individuals, allowing detailed analysis of both serologic and cellular immune memory to SARS-CoV-2 antigens.

**Figure 1.**
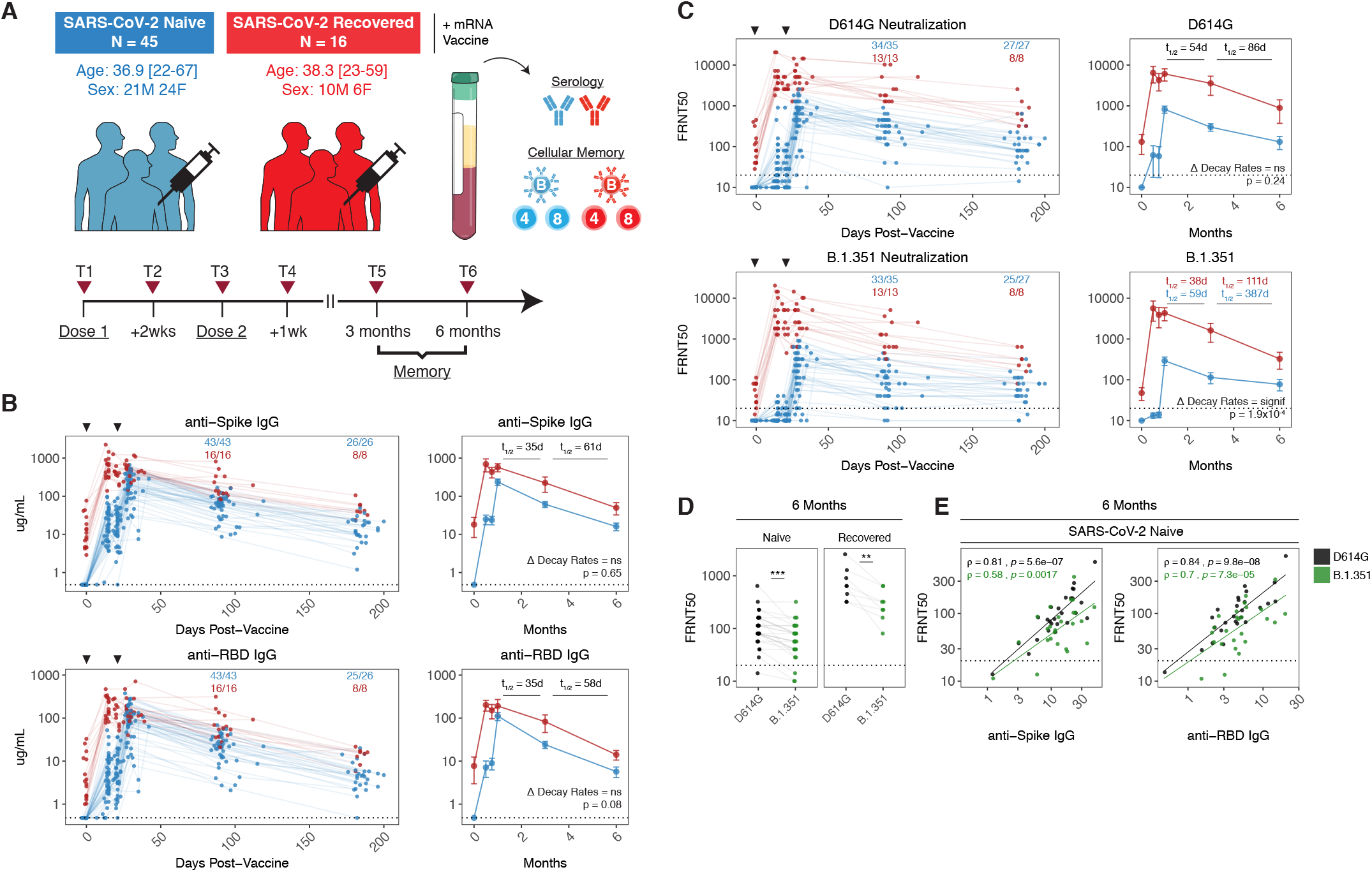
SARS-CoV-2 mRNA vaccines induce robust antibody responses. **A)** University of Pennsylvania COVID-19 vaccine study design and cohort summary statistics. **B)** anti-Spike and anti-RBD IgG concentrations over time in plasma samples from vaccinated individuals. **C)** Pseudovirus neutralization titers against wild-type D614G or B.1.351 variant Spike protein over time in plasma samples from vaccinated individuals. Data are represented as focus reduction neutralization titer 50% (FRNT50) values. **D)** Comparison of D614G and B.1.351 FRNT50 values at 6 months post-vaccination. Statistics were calculated using paired non-parametric Wilcoxon test **E)** Correlation between anti-Spike or anti-RBD IgG and neutralizing titers (D614G = black, B.1.351 = green; statistics were calculated using non-parametric Spearman rank correlation). Dotted lines indicate the limit of detection for the assay. For **B** and **C**, fractions above plots indicate the number of individuals above their individual baseline at memory timepoints. Decay rates were calculated using a piecewise linear mixed effects model with censoring. Decay rates were similar in SARS-CoV-2 naïve and recovered groups unless otherwise indicated. Blue and red values indicate comparisons within naïve or recovered groups.

### Antibody Responses to SARS-CoV-2 mRNA Vaccines

We first measured anti-Spike and anti-RBD binding antibody responses in plasma samples by enzyme linked immunosorbent assay (ELISA). As reported previously by our group and others, mRNA vaccines induced robust circulating antibody responses to the SARS-CoV-2 Spike protein and Spike RBD with distinct patterns of early response in SARS-CoV-2 naïve and recovered individuals (figure 1B) (*12, 29–31*). Peak levels of anti-Spike and anti-RBD IgG were observed 1 week after the second vaccine dose and subsequently declined over the course of the next 2 months with a half-life of ∼35 days (figure 1B), consistent with the dynamics of a typical immune response including a contraction of the vaccine-induced antibody secreting cell population. This decrease in antibody levels continued with a half-life of 58-61 days from 3-6 months post-vaccination (figure 1B). Of note, the calculated two-phase decay rates for anti-Spike and anti-RBD IgG were not significantly different between SARS-CoV-2 naïve and recovered vaccinees. Even after the decrease from peak antibody responses, all individuals had detectable anti-Spike IgG at 6 months.

To examine the functional quality of circulating antibodies, we used a neutralization assay with pseudotyped virus expressing either the wild-type Spike with the prevailing D614G mutation or the B.1.351 variant Spike (sequences listed in methods). We focused on B.1.351 neutralization as this variant has consistently shown the highest immune evasion among the current VOCs. In line with our binding antibody data, neutralizing titers for D614G and B.1.351 declined from peak levels after the second dose to 6 months for both SARS-CoV-2 naïve and recovered vaccinees (figure 1C). However, neutralizing titers displayed different decay kinetics, with slightly longer half-lives than binding antibody responses. Modelled decay rates for D614G neutralization were not significantly different between SARS-CoV-2 naïve and recovered vaccinees with a half-life of 86 days between 3-6 months post-vaccination (figure 1C). In contrast, a relative stabilization of neutralizing titers against the B.1.351 variant was observed between 3 and 6 months post-vaccination in individuals without a prior SARS-CoV-2 infection with a half-life of 387 days, compared to 111 days in SARS-CoV-2 recovered subjects (figure 1C). We next compared neutralizing titers to D614G and B.1.351 at 6 months post-vaccination. Despite a significant loss of neutralizing ability against B.1.351 relative to D614G (figure 1D), 24/27 SARS-CoV-2 naïve and 8/8 SARS-CoV-2 recovered individuals still had neutralizing titers above the limit of detection at 6 months post-vaccination. Finally, cross-sectional analysis of 6-month antibody responses also demonstrated that binding antibodies remained highly correlated with neutralizing titers (figure 1E), indicating that Spike- and RBD-specific antibody responses retain their functional characteristics and neutralizing capacity over time.

### Memory B Cell Responses to SARS-CoV-2 mRNA Vaccines

In addition to antibodies, we measured the frequencies of SARS-CoV-2 Spike- and RBD-specific memory B cells in peripheral blood using a flow cytometric assay. Antigen specificity was determined based on binding to fluorescent SARS-CoV-2 Spike and RBD probes (figure 2A-B). Influenza hemagglutinin (HA) from the 2019 flu vaccine season was also included as a historical antigen control. Full gating strategies are provided in **figure S1**.

**Figure 2.**
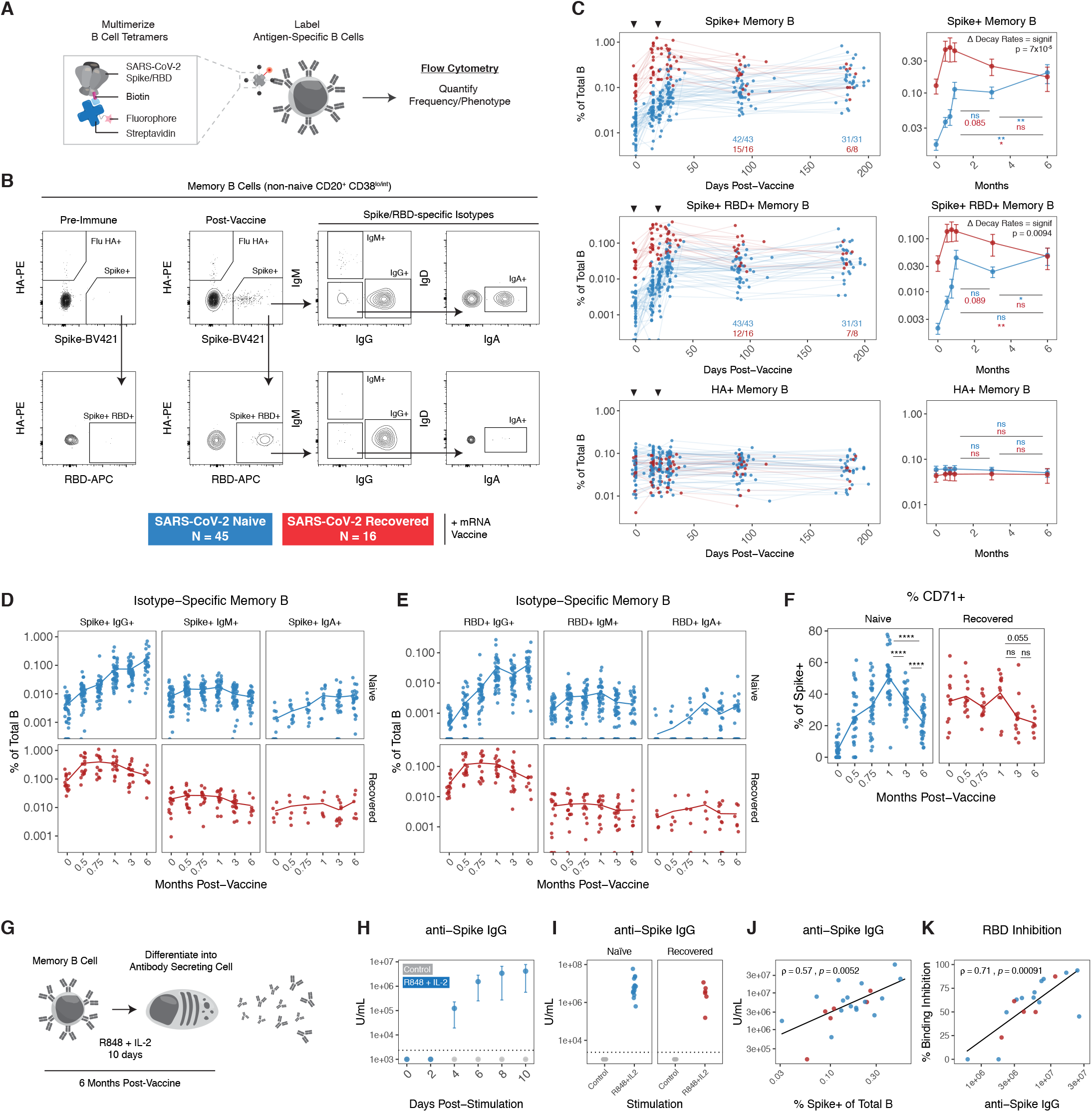
SARS-CoV-2 mRNA vaccines generate durable and functional memory B cell responses. **A)** Experimental design and **B)** gating strategy for quantifying the frequency and phenotype of SARS-CoV-2-specific memory B cells by flow cytometry. Antigen specificity was determined based on binding to fluorophore-labeled Spike, RBD, and influenza HA tetramers. **C)** Frequencies of SARS-CoV-2 Spike+, Spike+ RBD+, and influenza HA+ memory B cells over time in PBMC samples from vaccinated individuals. Data are represented as a percentage of total B cells and fractions above plots indicate the number of individuals above their individual baseline at memory timepoints. Decay rates were calculated using a piecewise linear mixed effects model with censoring. **D)** Frequency of isotype-specific Spike+ and **E)** Spike+ RBD+ memory B cells over time. **F)** Percent CD71+ of total Spike+ memory B cells over time. **G)** Experimental design for differentiation of memory B cells into antibody secreting cells. **H)** anti-Spike IgG levels in culture supernatants over time from PBMCs stimulated with PBS control or R848 + IL-2 (n=4). **I)** anti-Spike IgG levels in culture supernatants after 10 days of stimulation **J)** Correlation of Spike+ memory B cell frequencies by flow cytometry with anti-Spike IgG levels from *in vitro* stimulation. **K)** Correlation of anti-Spike IgG levels from *in vitro* stimulation with RBD-binding inhibition. For **J** and **K**, correlations were calculated using non-parametric Spearman rank correlation. Statistics were calculated using unpaired non-parametric Wilcoxon test with Holm correction for multiple comparisons. Blue and red values indicate comparisons within naïve or recovered groups.

SARS-CoV-2-specific memory B cells were detectable in all previously uninfected individuals after 2 vaccine doses and remained stable as a percentage of total B cells from 1-3 months post-vaccination (figure 2C). All SARS-CoV-2 recovered individuals in our study had detectable antigen-specific memory B cells at pre-vaccination baseline and these pre-existing memory B cells were significantly boosted by the first vaccine dose, with little change after the second vaccine dose (figure 2C). No changes were observed in influenza HA+ memory B cells after SARS-CoV-2 vaccination for either group.

Longitudinal analysis revealed a continued increase in the frequency of Spike+ and Spike+ RBD+ memory B cells from 3-6 months post-vaccination in SARS-CoV-2 naïve individuals, whereas the frequency of these antigen-specific memory B cells in SARS-CoV-2 recovered subjects continued to decline from peak levels (figure 2C). One possible explanation for the observed increase in frequency of vaccine-induced memory B cells over time in SARS-CoV-2 naïve vaccinees is prolonged germinal center activity, resulting in continued export of memory B cells. Indeed, antigen-specific germinal center B cells have been documented in axillary lymph nodes at 15 weeks post-mRNA vaccination in SARS-CoV-2 naïve subjects (*11*), though germinal center dynamics in vaccinees with prior immunity to SARS-CoV-2 remain to be defined. SARS-CoV-2 recovered individuals had consistently higher frequencies of antigen-specific memory B cells up to 3 months post-vaccination (figure 2C). However, due to distinct trajectories, both SARS-CoV-2 naïve and SARS-CoV-2 recovered individuals had similar frequencies of Spike+ and Spike+ RBD+ memory B cells at 6 months post-vaccination (figure 2C), perhaps reflecting some upper limit to the frequencies of antigen-specific memory B cells that can be maintained long-term.

We next investigated the phenotype of mRNA vaccine-induced memory B cells. Analysis of immunoglobulin isotypes in SARS-CoV-2 naïve vaccinees revealed a steady increase in IgG+ memory B cells over time (figure 2D-E, **figure S2A-C**), indicating ongoing class-switching. By contrast, IgM+ cells were most abundant at baseline and early post-vaccination timepoints. IgM+ and IgA+ memory B cells represented a minor fraction of the overall response in the blood at later timepoints. In SARS-CoV-2 recovered vaccinees, the majority of Spike+ and Spike+ RBD+ memory B cells were IgG+ at baseline, and the fraction of IgG+ cells was further boosted by vaccination (figure 2D-E**, figure S2A-C**). Moreover, we assessed the activation status of antigen-specific memory B cells by CD71 expression (*32*). The percent of Spike+ memory B cells expressing CD71 increased over the course of the 2-dose vaccine regimen in SARS-CoV-2 naïve individuals, peaking at 1 week after the second vaccine dose (figure 2F). The percent of CD71+ antigen-specific memory B cells then steadily declined through the 6-month timepoint, indicating a transition towards a population of mature resting memory B cells. A similar decrease in CD71 expression was observed from 1-6 months post-vaccination in SARS-CoV-2 recovered individuals (figure 2F).

Given the robust generation of Spike- and RBD-binding memory B cells, we next tested whether vaccine-induced memory B cells could produce functional antibodies upon reactivation. This reactivation-induced antibody production may be especially relevant in the setting of antigen re-encounter, either through exposure to live virus or an additional vaccine dose. To this end, we established an *in vitro* culture system to differentiate memory B cells into antibody secreting cells (*33*). PBMC samples from vaccinated individuals at the 6-month timepoint were cultured with a combination of R848 and IL-2 and culture supernatants were collected to measure antibody levels and function (figure 2G). Robust anti-Spike IgG was detected in supernatants as early as 4 days post-stimulation (figure 2H), indicating that memory B cells can act as a rapid source of secondary antibody production. All 6-month samples generated significant levels of anti-Spike IgG in this assay compared to unstimulated controls (figure 2I). This *in vitro* anti-Spike IgG production also correlated strongly with the frequency of Spike+ memory B cells detected by flow cytometry (figure 2J), supporting the functional relevance of quantifying antigen-specific memory B cells in the blood. We further tested the function of memory B cell-derived antibodies from culture supernatants using an ELISA-based ACE2-binding inhibition assay as a surrogate of neutralization capacity. Indeed, ACE2-binding inhibition activity correlated strongly with the levels of anti-Spike IgG antibody in culture supernatant (figure 2K). Taken together, these data demonstrate that mRNA vaccines induce a population of memory B cells that are durable for at least at 6 months after vaccination and that are capable of rapidly producing functional antibodies against SARS-CoV-2 upon stimulation.

### Memory B Cell Responses to Major Variants of Concern (VOCs)

A major question regarding anti-SARS-CoV-2 immunity is the extent to which memory B cells induced by current mRNA vaccines can bind different regions of the Spike protein, and how these responses may be affected by emerging VOCs. We therefore examined vaccine-induced memory B cell responses targeting different components of the Spike protein as well as current VOCs using an expanded antigen probe panel. Specifically, we designed B cell tetramers for 8 SARS-CoV-2 antigens, including full-length Spike, N-terminal domain (NTD), multiple variant RBDs (wild-type, B.1.1.7, B.1.351, and B.1.617.2), and the S2 domain (figure 3A-B). Spike-specific memory B cells were defined based on a multiple-discrimination approach, with binding to full-length Spike plus one or more additional probes. This strategy also allowed us to identify memory B cells that cross-bind all variant RBDs (RBD++++). SARS-CoV-2 nucleocapsid was used as a vaccine-irrelevant antigen (but one for which SARS-CoV-2 immune subjects had detectable pre-existing immunity; **figure S3A-B**). Full gating strategies are provided in **figure S1**. We also leveraged a separate cohort of healthcare workers (HCW) who had mild COVID-19 and were sampled longitudinally after a positive serology test to compare vaccine-induced responses with natural infection (*34*).

**Figure 3.**
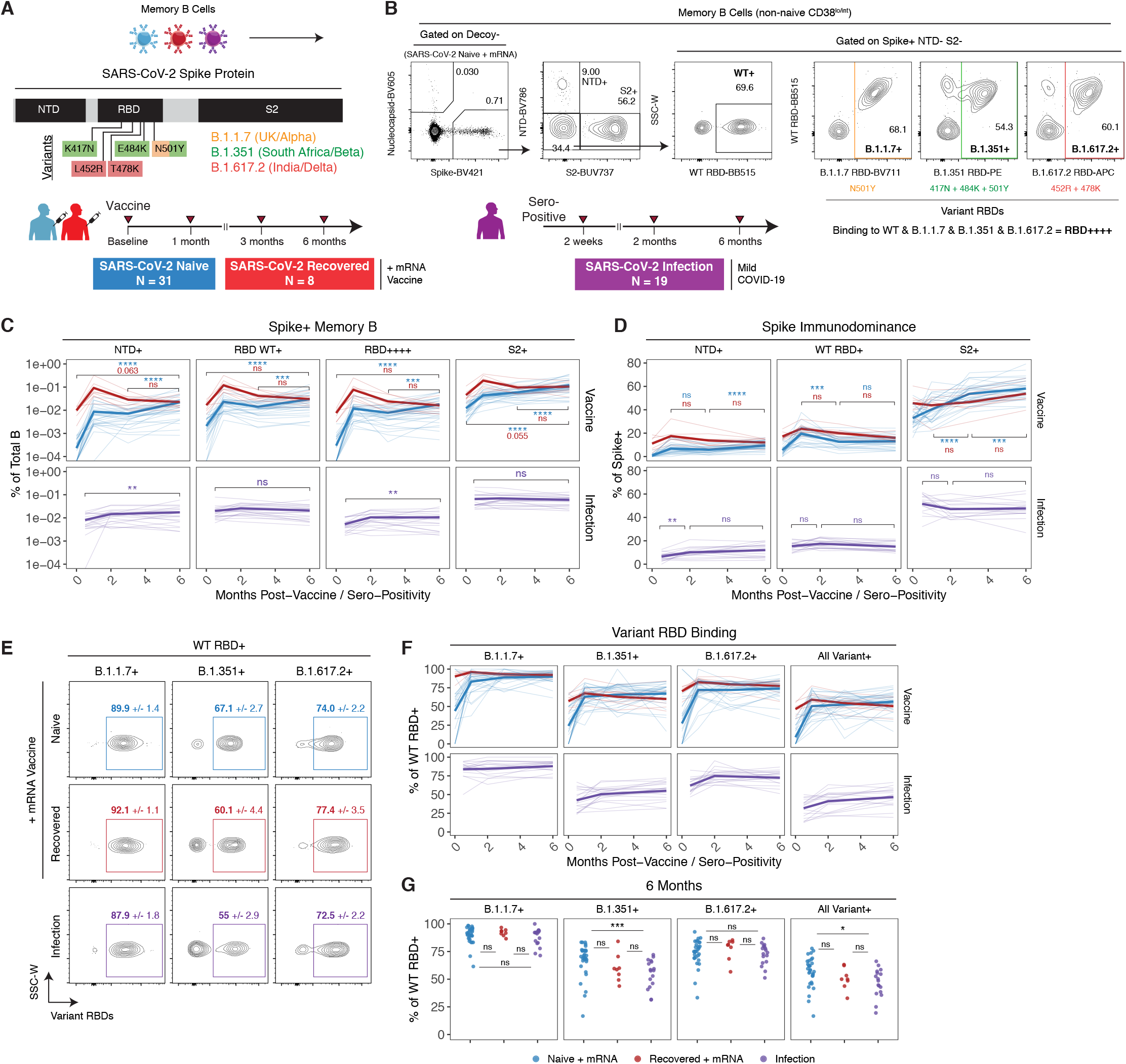
Memory B cells induced by mRNA vaccination are cross-reactive to SARS-CoV-2 variants of concern and increase in frequency over time. **A)** Experimental design and **B)** gating strategy for quantifying the frequency and phenotype of Spike component and variant-specific memory B cells by flow cytometry. Specific mutations in B.1.1.7, B.1.351, or B.1.617.2 variant RBDs are indicated. **C)** Frequencies of Spike+ NTD+, Spike+ WT RBD+, Spike+ RBD++++ (all variant binding), and Spike+ S2+ memory B cells over time in PBMC samples from vaccinated or convalescent individuals. Data are represented as a percentage of total B cells. **D)** Percent NTD+, RBD+, or S2+ of total Spike+ memory B cells over time. **E)** Representative plots of variant RBD cross-binding gated on Spike+ WT RBD+ cells in vaccinated or convalescent individuals. Mean and standard error values at the 6-month timepoint are indicated. **F)** Percent B.1.1.7+, B.1.351+, or B.1.617.2+ of WT RBD+ memory B cells over time. **G)** Cross-sectional analysis of variant binding as a percentage of WT RBD+ memory B cells at 6 months post-vaccination/seropositivity. Statistics were calculated using paired non-parametric Wilcoxon test with Holm correction for multiple comparisons. Blue, red, and purple values indicate comparisons within naïve, recovered, or natural infection groups.

mRNA vaccination induced robust memory B cell responses to all SARS-CoV-2 Spike antigens, and the frequency of these memory B cells increased from 3-6 months post-vaccination in SARS-CoV-2 naïve individuals (figure 3C). In the mild natural infection HCW cohort, a gradual increase in the frequency of Spike+ NTD+ and Spike+ RBD++++ memory B cells was also observed from 2 weeks to 6 months post-seropositive test. Cross-sectional analysis at 6 months post-vaccination or sero-positivity revealed similar antigen-specific memory B cell frequencies between all groups (**figure S3B**), suggesting that both vaccination and infection can induce durable memory B cell populations.

As our panel included probes covering much of the Spike protein, including NTD, RBD, and S2, we also examined immunodominance patterns and how B cell immunodominance to Spike changed over time. In previously uninfected individuals, ∼30-40% of Spike-binding memory B cells co-bound S2 at pre-vaccine baseline (figure 3D). Previous work has shown that the S2 domain of SARS-CoV-2 Spike is more conserved with other coronaviruses and it is likely that S2-binding memory B cells detected at baseline reflect cross-reactivity to these commonly circulating coronaviruses (*35, 36*). mRNA vaccination induced robust populations of S2-specific memory B cells in SARS-CoV-2 naïve vaccinees, with S2+ cells accounting for ∼40-80% of the total Spike-specific memory B cell population at 6 months. Although the overall frequency of NTD+ and RBD+ memory B cells increased over time, they were comparatively less immunodominant than S2 as a percentage of total Spike+ memory B cells (figure 3C-D). mRNA vaccination induced a gradual increase in NTD-specificity over time in SARS-CoV-2 naïve individuals, whereas RBD-specificity as a percent of Spike+ memory B cells appeared to have a more prominent peak 1 week after the second vaccine dose and then stabilized from 3-6 months post-vaccination (figure 3D). When SARS-CoV-2 recovered subjects were boosted by mRNA vaccination, a similar immunodominance pattern was observed with S2-specificity representing most of the total Spike response (figure 3D). Overall, vaccination did not change the pre-existing immunodominance of NTD, RBD, and S2 over time in this group, suggesting that boosting previous immunity with mRNA vaccine in SARS-CoV-2 recovered individuals provides little long-term benefit to the frequency or proportions of memory B cells binding different SARS-CoV-2 Spike antigens. In the context of natural infection only, we similarly found that NTD, RBD and S2 immunodominance remain relatively constant from early convalescence through late memory (figure 3D).

We next examined memory B cell binding to B.1.1.7 (Alpha), B.1.351 (Beta), and B.1.617.2 (Delta) variant RBDs relative to WT RBD (figure 3E-F). All RBD probes were used at the same concentration to facilitate direct comparisons and specific point mutations are shown in figure 3A-B. Variant-binding memory B cells were detectable in all SARS-CoV-2 naïve individuals after 2 vaccine doses and increased slightly as a percentage of WT RBD+ cells from 1-6 months post vaccination (figure 3F). By contrast, vaccination induced a transient increase in variant cross-binding as a percent of WT RBD+ memory B cells in SARS-CoV-2 recovered individuals, but this effect was lost over time and variant cross-binding returned to pre-vaccination levels at 6 months (figure 3F). In convalescent individuals who recovered from a mild infection, there was a gradual increase in variant cross-binding over time (figure 3F). Class-switching to an IgG dominated response was also observed in all groups (**figure S3C-D**). Of note, the variants and corresponding mutations tested in our panel had different magnitudes of effect (figure 3E-F). B.1.1.7 RBD with a single N501Y mutation had relatively little change in binding compared to WT RBD. As expected, B.1.351 RBD resulted in a more substantial loss of cross-binding. B.1.617.2 RBD had an intermediate effect on binding.

Cross-sectional analysis of variant-binding at the 6-month timepoint also revealed two major findings. First, all vaccinated individuals in our study maintained variant-specific memory B cells for at least 6 months, with an average of >50% of WT RBD+ memory B cells also cross-binding all 3 major variants of concern (figure 3G). Second, mRNA vaccination in SARS-CoV-2 naïve individuals induced a stronger response to B.1.351 than natural infection alone (figure 3G). One possible explanation for this difference is the immunogen itself. Vaccinated individuals mount a primary response to the mRNA-encoded prefusion stabilized Spike trimer, potentially allowing increased recruitment and/or selection of specific clones that can bind conserved regions of RBD (*37, 38*). In contrast, convalescent individuals were primed against native, non-stabilized Spike protein. Taken together, our data indicate robust B cell memory to multiple components of the Spike protein as well as currently described VOCs that continues to evolve and increase in frequency over time.

### Clonal Evolution of Variant-Specific Memory B Cells

We next asked what differences may underly variant-binding versus non-binding properties of vaccine-induced memory B cells. Here, we focused on the B.1.351 variant RBD containing the K417N, E484K, and N501Y mutations as this variant resulted in the greatest loss of binding relative to WT RBD. We designed a sorting panel to identify 3 populations of memory B cells with different antigen-binding specificities: 1) memory B cells that bind full-length Spike but not RBD, 2) memory B cells that bind full-length Spike and WT RBD but not B.1.351 variant RBD, and 3) memory B cells that bind full-length Spike and cross-bind both WT and B.1.351 variant RBD. Naïve B cells were also sorted as a control. These populations were isolated from 8 SARS-CoV-2 naïve and 4 SARS-CoV-2 recovered individuals at 3-4 months post-vaccination (figure 4A**, figure S4A**). Consistent with our previous data, between 50-80% of WT RBD+ cells co-bound B.1.351 variant RBD (figure 4B), indicating that a majority of RBD epitopes in the response are shared by the WT and mutant RBDs.

**Figure 4.**
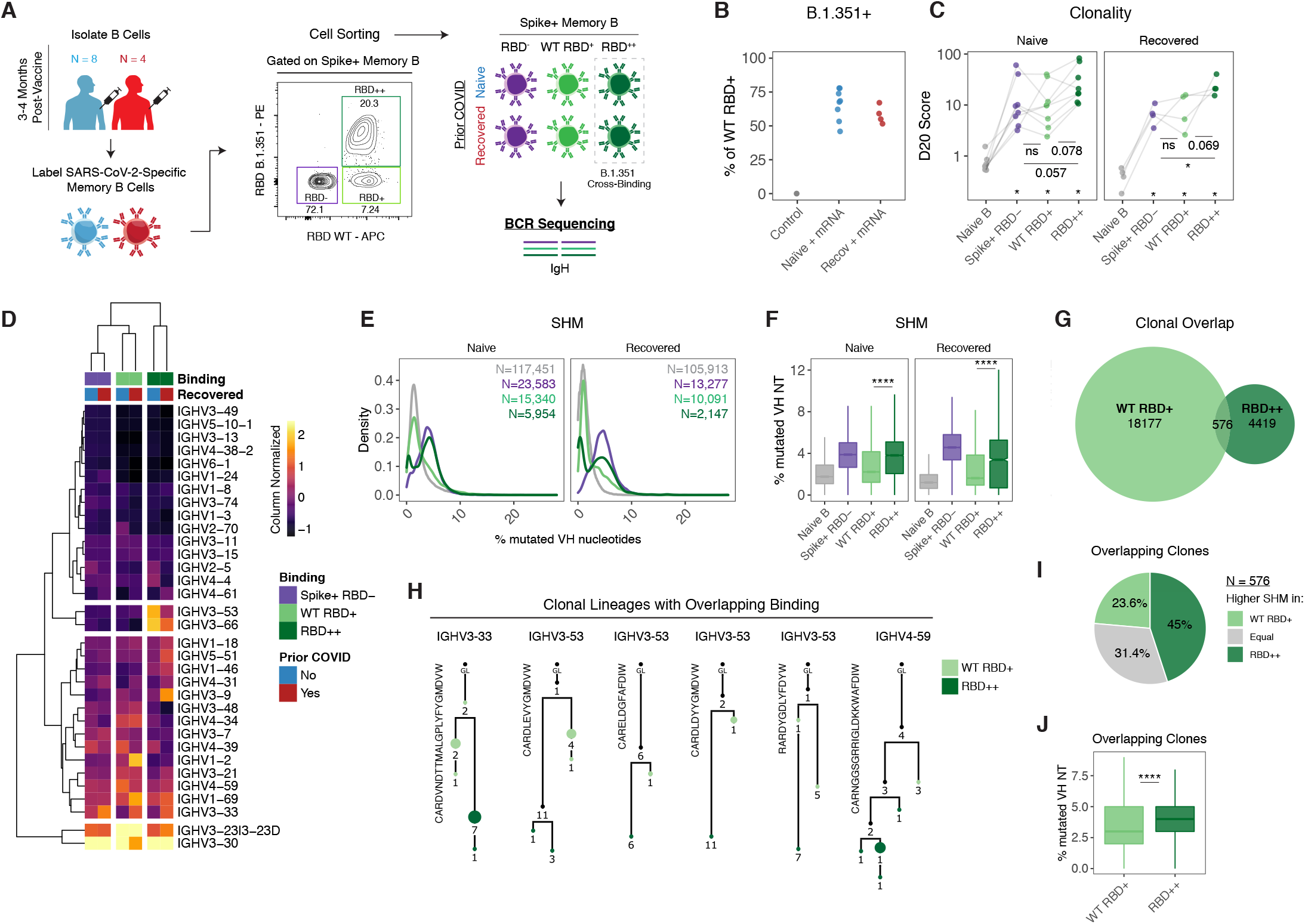
Variant-binding memory B cell clones use distinct VH genes and evolve through somatic hypermutation. **A)** Experimental design for sorting and sequencing SARS-CoV-2-specific memory B cells. **B)** Frequency of RBD++ (cross-binding) memory B cells as a percentage of total RBD+ cells. **C)** Percentage of sequence copies occupied by the top 20 ranked clones (D20) across naïve B cells and different antigen-binding memory B cell populations. **D)** Heatmap and hierarchical clustering of VH gene usage frequencies in memory B cell clones across different antigen-binding memory B cell populations. Data are represented as column normalized values. **E)** Somatic hypermutation (SHM) distributions and **F)** boxplots of individual clones across naïve B cells and different antigen-binding memory B cell populations. Data are represented as the percent of mutated VH nucleotides. Number of clones sampled for each population is indicated. **G)** Venn diagram of clonal lineages that are shared between WT RBD and RBD cross-binding (RBD++) populations. Data were filtered based on larger clones with ≥ 50% mean copy number frequency in each sequencing library. **H)** Example lineage trees of clones with overlapping binding to WT and B.1.351 variant RBD. VH genes and CDR3 sequences are indicated. Numbers refer to mutations compared to the preceding vertical node. Colors indicate binding specificity, black dots indicate inferred nodes, and size is proportional to sequence copy number; GL = germline sequence. **I)** Classification of SHM within overlapping clones. Each clone was defined as having higher (or equal) SHM in WT RBD binders or RBD++ cross-binders based on average levels of SHM for all WT RBD vs. RBD++ sequence variant copies within each lineage. **J)** SHM levels within overlapping clones. Data are represented as the percent of mutated VH nucleotides for WT RBD and RBD++ sequence copies. Statistics were calculated using paired non-parametric Wilcoxon test, with Holm correction for multiple comparisons in **C**.

To gain insight into the clonal composition of the different spike and/or RBD-binding B cell populations, IgH rearrangements were amplified from the sorted populations (N=48 total) and related sequences were grouped into clones (N=348,346 clones, **table S2**). We analyzed the contribution of the top copy number clones to the overall repertoire as measured by the D20 index. The D20 index ranged from less than 1% for naïve B cells (which is expected for a diverse, non-clonally expanded population) to greater than 20% for some of the antigen-binding populations (figure 4C). Clones that cross-bound both WT and B.1.351 RBD had the highest D20 scores, indicating greater clonal expansion and/or lower diversity compared to the other antigen-binding populations (figure 4C). Although the clonality of antigen-binding memory B cell populations was not significantly different after vaccination based on prior immunity, there was considerable heterogeneity in clonal expansion across individuals, with multiple SARS-CoV-2 naïve subjects showing higher D20 scores compared to those who were previously infected and then boosted by mRNA vaccine.

We further analyzed IGHV gene usage across the different antigen-binding memory B cell populations. Hierarchical clustering revealed that VH gene profiles were overall similar in vaccinated individuals regardless of prior SARS-CoV-2 infection status (figure 4D**, figure S4B**), indicating that both vaccination and infection boosted by vaccination can recruit similar clones into the response. Rather, IGHV gene usage largely clustered based on the antigen-binding phenotype, with increased usage of VH3-53 and VH3-66 in RBD cross-binding clones (figure 4D**, figure S4B**). Of note, both of these IGHV genes are known to be enriched in spike-binding B cells (*39, 40*). These differences in IGHV gene usage between WT only and variant cross-binding phenotype suggest that these cells may derive, at least partially, from different B cell clones that were independently recruited into the vaccine response.

Analysis of somatic hypermutation across different antigen-binding populations also revealed several major findings. As expected, SARS-CoV-2-specific memory B cell clones had significantly more VH nucleotide mutations compared to naïve B cell clones (figure 4E-F**, figure S4C**). However, there were clear differences in SHM between the different antigen-binding populations. Spike+, RBD non-binding memory B cells (which include NTD-and S2-binding populations) had high SHM (figure 4E-F**, figure S4C**), consistent with germinal center-dependent responses as well as possible recall responses of pre-existing cross-reactive S2 clones. Significantly higher levels of SHM were also observed in variant RBD cross-binding clones compared to WT RBD only clones (figure 4E-F**, figure S4C**). Notably, boosting of natural immunity by mRNA vaccination in SARS-CoV-2 recovered donors did not appear to produce higher SHM in memory B cell clones for any of the different antigen-binding specificities examined (figure 4F).

To determine if variant cross-binding clones could evolve from WT RBD-binding clones, we next investigated if there was any clonal overlap between these populations. For clonal overlap analysis, we focused on larger clones (defined as having copy numbers at or above 50% of the mean copy number frequency within each sequencing library) (*41*), as larger clones are more readily sampled at both the clonal and subclonal levels. Among such larger clones, 2.5% had sequence variants that were isolated from both WT RBD and cross-binding populations (figure 4G**, figure S4D**). Lineage analysis revealed that WT and variant cross-binding sequence variants localized on separate branches (representative lineages shown in figure 4H), indicating that the shift in antigen-reactivity was not due to contamination of the sorted populations (in which case sequence variants localize to the same nodes). Next, to determine if cross-binding activity arose from WT binding or vice versa, we used SHM as a molecular clock and counted the fraction of overlapping clonal lineages in which variant binding had higher, lower, or equivalent levels of SHM to WT RBD-binding variants. Consistent with the overall SHM data, this analysis of overlapping clones also revealed higher levels of SHM in the variant binding sequences compared to WT only binding sequences (figure 4I-J).

Taken together, these data indicate that mRNA vaccine-induced memory B cells that bind variant RBDs have higher SHM compared to clones that only bind WT RBD. Moreover, the clonal relationships between WT-only and cross-binding RBD-specific memory B cells suggest that variant binding capacity can evolve from clones that initially bound to WT RBD. Ongoing evolution and selection of these clones could therefore facilitate cross-protection against different VOCs. This finding is consistent with earlier work suggesting that SHM and affinity maturation are important for the acquisition of broader neutralization activity of RBD-binding antibodies that are formed in response to SARS-CoV-2 infection (*42, 43*). Future work is necessary to determine how additional antigen exposure through booster vaccination, environmental virus exposure, or overt infection may impact additional affinity maturation towards improved variant-binding.

### Memory CD4+ and CD8+ T Cell Responses to SARS-CoV-2 mRNA Vaccines

In addition to antibodies and memory B cells, memory T cells can contribute to protection upon re-exposure to virus. Memory T cell responses have also been shown to be less affected by variants of concern than humoral immune responses (*17, 44*). To determine whether mRNA vaccination induced durable antigen-specific memory T cell responses, we performed a flow cytometric analysis using an activation induced marker (AIM) assay. PBMCs were stimulated with peptide megapools containing optimized Spike epitopes (*45, 46*). Antigen-specific responses were quantified based on the frequency of AIM^+^ non-naïve T cells in stimulated samples above background frequencies in paired unstimulated controls (figure 5A-B) (*15*). Full gating strategies are provided in **figure S5**. Antigen-specific CD4+ T cells were defined based on co-expression of CD40L and CD200. Antigen-specific CD8+ T cells were defined based on expression of 4 of 5 total activation markers as described previously (*15*).

**Figure 5.**
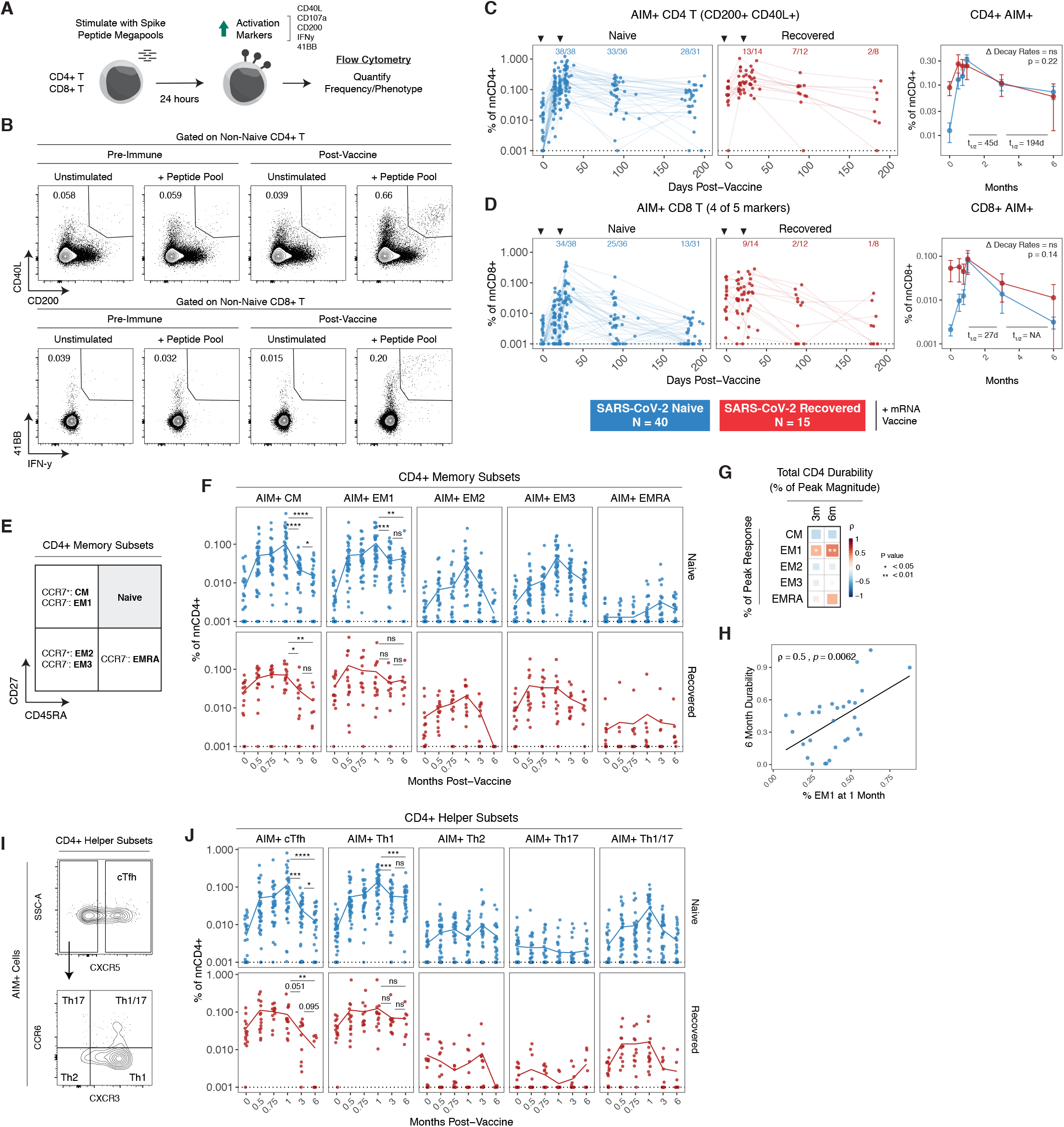
SARS-CoV-2 mRNA vaccines generate durable memory T cell responses. **A)** Experimental design and **B)** gating strategy for quantifying the frequency of SARS-CoV-2-specific CD4+ and CD8+ T cells by AIM assay. For CD4+ T cells, antigen specificity was defined based on co-expression of CD40L and CD200. For CD8+ T cells, antigen specificity was defined based on expression of at least 4/5 activation markers. **C)** Frequencies of AIM+ CD4+ T and **D)** AIM+ CD8+ T cells over time in PBMC samples from vaccinated individuals. Data were background subtracted using a paired unstimulated control for each timepoint and are represented as a percentage of non-naïve CD4+ or CD8+ T cells. Fractions above plots indicate the number of individuals above their individual baseline at memory timepoints. Decay rates were calculated using a piecewise linear mixed effects model with censoring. Decay rates were similar in SARS-CoV-2 naïve and recovered groups. **E)** AIM+ CD4+ T cell memory subsets were identified based on surface expression of CD45RA and CD27, followed by CCR7. **F)** Frequencies of AIM+ CD4+ T cell memory subsets over time. **G)** Correlation matrix of memory subset skewing at peak (1 month) response with total AIM+ CD4+ T cell durability at 3 and 6 months. Durability was measured as the percent of peak response maintained at memory timepoints for each individual. **H)** Correlation between percent of EM1 cells at peak response with 6-month durability. **I)** AIM+ CD4+ T helper subsets were defined based on chemokine receptor expression. **J)** Frequencies of AIM+ CD4+ T helper subsets over time. Dotted lines indicate the limit of detection for the assay. Statistics were calculated using unpaired non-parametric Wilcoxon test with Holm correction for multiple comparisons. Correlations were calculated using non-parametric Spearman rank correlation.

Consistent with recent studies, SARS-CoV-2 mRNA vaccination efficiently primed antigen-specific CD4^+^ T cells and CD8^+^ T cells (figure 5C-D) (*16–18*). All individuals in our cohort, regardless of prior infection with SARS-CoV-2, had detectable CD4^+^ T cell responses above their individual baseline one week following the second vaccine dose (figure 5C). Most SARS-CoV-2 naïve individuals also generated detectable CD8^+^ T cell responses after the second dose (figure 5D). In contrast, vaccination did little to further boost pre-vaccination antigen-specific CD8^+^ T cell frequencies in SARS-CoV-2 recovered individuals (figure 5D). A marked contraction phase was observed from peak responses to 3-months post-vaccination, with a half-life of 45 days for CD4^+^ T cells and 27 days for CD8^+^ T cells (figure 5C-D). These kinetics are consistent with a typical T cell response after the effector phase (*47*). After this initial contraction, antigen-specific memory CD4+ T cell frequencies stabilized from 3-6 months post-vaccination with a half-life of 194 days, whereas CD8+ T cells continued to decline. Overall, 28/31 SARS-CoV-2 naïve individuals had vaccine-induced antigen-specific CD4^+^ T cell responses at 6 months post-vaccination above pre-vaccination baseline levels, and 13/31 had detectable CD8^+^ T cell responses above baseline (figure 5C-D). In SARS-CoV-2 recovered subjects, mRNA vaccination had only a modest impact on T cell responses and did not improve the overall magnitude of long-term antigen-specific CD4+ or CD8+ T cell memory (figure 5C-D). Taken together, these data indicate that mRNA vaccination generates durable SARS-CoV-2-specific CD4+ T cell memory in individuals who were not previously infected with SARS-CoV-2 and only transiently boosts these responses in SARS-CoV-2 recovered individuals.

Antigen-specific T cells can further be classified into different memory subsets using cell surface markers (figure 5E). Peak CD4+ T cell responses following SARS-CoV-2 mRNA vaccination were composed of predominantly central memory (CM; CD45RA-CD27+ CCR7+) and effector memory 1 (EM1; CD45RA-CD27+ CCR7-) cells in both SARS-CoV-2 naïve and recovered individuals (figure 5F) (*15*). During contraction from peak responses, antigen-specific CCR7^+^ CM cells were largely lost from circulation, whereas antigen-specific CCR7^-^ EM1 cells stabilized in frequency from 3-6 months post-vaccination. Moreover, the percentage of the peak CD4+ response that was EM1 cells, but not other memory subsets, was significantly associated with the durability of the overall CD4+ T cell response at 3 and 6 months post-vaccination (figure 5G-H), suggesting that EM1s are long-lived memory CD4+ T cells and that early skewing towards an EM1 phenotype contributes to durable memory T cell responses. Although our AIM assay allows detection of low-frequency memory CD8^+^ T cell responses for overall quantification, reliable subsetting of antigen-specific CD8^+^ T cells at memory timepoints was not feasible due to the low number of events.

mRNA vaccination also preferentially induced antigen-specific CD4^+^ cTfh and Th1 helper cells in both SARS-CoV-2 naïve and recovered individuals, whereas Th2, Th17, and Th1/17 cells were generated at a lower level (figure 5I). Although the overall frequency of antigen-specific CD4^+^ T cells stabilized from 3-6 months post-vaccination, cTfh and Th1 cells had distinct trajectories. Specifically, cTfh cells declined more rapidly than Th1 cells both during the initial contraction phase and from 3-6 months post-vaccination (figure 5J). In contrast, Spike-specific Th1 cells did not decline from 3-6 months post-vaccination. While cTfh cells may be important in the early stages of vaccine response, these data indicate that the durable component of the memory CD4+ T cell response at 6 months post-vaccine is largely composed of Th1 cells, and boosting of pre-existing immunity with mRNA vaccine does not change the magnitude or subset composition of the CD4+ memory T cell response.

### Integrated Analysis of Immune Components and Vaccine-Induced Memory to SARS-CoV-2

A goal of this study was to assess the development of multiple components of antigen-specific immune memory over time in the same individuals following SARS-CoV-2 mRNA vaccination. This dataset allowed us to integrate longitudinal antibody, memory B cell, and memory T cell responses to construct an immunological landscape of SARS-CoV-2 mRNA vaccination in previously uninfected subjects. To this end, we applied uniform manifold approximation and projection (UMAP) to visualize the trajectory of vaccine-induced adaptive immunity over time. This analysis revealed a continued evolution of the overall immune response in SARS-CoV-2 naïve subjects after mRNA vaccination with different timepoints occupying largely non-overlapping UMAP space (figure 6A). Projection of individual immune components onto the UMAP space revealed that primary vaccination was largely defined by rapid induction of CD4+ T cell immunity (figure 6B). The second vaccine dose induced peak antibody, CD4+ T cell, and CD8+ T cell responses. Antibodies and CD4+ T cells then remained durable through later memory timepoints, coinciding with a trajectory shift towards peak memory B cell responses. Notably, all 6-month samples clustered away from pre-immune baseline samples (figure 6A), highlighting the durable multi-component immune memory induced by mRNA vaccination. At 6 months, we observed some heterogeneity in the immune landscape. This heterogeneity may be partially driven by a significant negative correlation between age and anti-Spike IgG (**figure S6A-B**). Sex did not appear to have any association with the overall antigen-specific response to mRNA vaccination (**figure S6C**).

**Figure 6.**
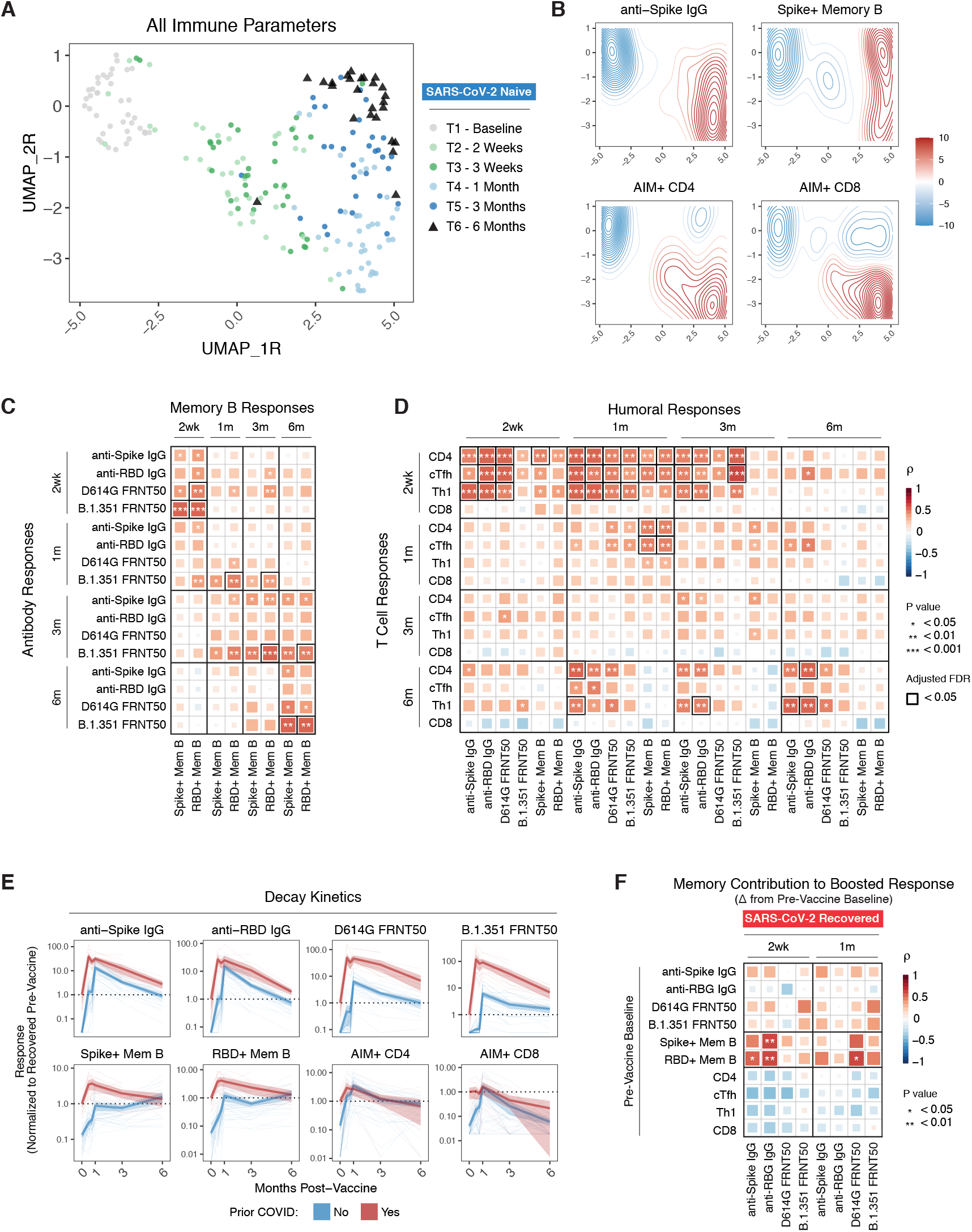
Immune trajectories and relationships in response to SARS-CoV-2 mRNA vaccination. **A)** UMAP of 12 antigen-specific parameters of antibody, memory B, and memory T cell responses to mRNA vaccination in SARS-CoV-2 naïve subjects. Data points represent individual participants and are colored by timepoint relative to primary vaccine. Triangles indicate 6-month samples. **B)** Kernel density plots of anti-Spike IgG, Spike+ memory B, AIM CD4+, and AIM+ CD8+ T cells. Red contours represent areas of UMAP space that are enriched for specific immune components. **C)** Correlation matrix of antibody and memory B cell responses over time in SARS-CoV-2 naïve subjects. **D)** Correlation matrix of T cell and humoral responses over time in SARS-CoV-2 naïve subjects. **E)** Decay kinetics of antibody, memory B cell, and memory T cell parameters over time in SARS-CoV-2 naïve and recovered vaccinees. Data are normalized to pre-vaccine levels in SARS-CoV-2 recovered individuals to evaluate the effect of boosting pre-existing immunity. **F)** Correlation matrix of baseline memory components with antibody recall responses after vaccination in SARS-CoV-2 recovered individuals. Recall responses were calculated as the difference between post-vaccination levels and pre-vaccine baseline. All statistics were calculated using non-parametric Spearman rank correlation. Black boxes indicate significant values after FDR correction.

A related question is how different antigen-specific mRNA vaccine-induced immune components interact with each other over time. Antibody responses after the first or second dose did not correlate with the magnitude of B cell memory at 6 months (figure 6C). However, at 3-and 6-months post-vaccination antibodies were significantly associated with contemporaneous memory B cell responses, an effect most prominent for B.1.351 neutralizing titers (figure 6C). Given the role of Tfh cells in generating efficient humoral immunity, we next investigated the relationship between antigen-specific T cells and humoral responses. CD4+ T cell responses, especially cTfh responses, as early as 2 weeks after the first dose of mRNA vaccine were positively correlated with antibody responses after the second vaccine dose, and this correlation persisted through 6 months (figure 6D**, figure S6D**). This observation suggests that rapid mobilization of CD4+ T cell responses by the first mRNA vaccine dose has a lasting effect on humoral immunity. Like memory B cells, the magnitude of CD4+ T cell responses at 6 months was also correlated with antibodies at 6 months (figure 6D), suggesting that antibody levels may provide a useful (though incomplete) proxy for the magnitude of memory B and CD4+ T cell responses at 6 months post-vaccination. Taken together, these data identify key temporal relationships between different branches of the human immune response that are associated with long-term immune memory after mRNA vaccination.

Next, we investigated if the magnitude of peak responses after the second vaccine dose in SARS-CoV-2 naïve subjects was predictive of memory responses at 3 and 6 months. Indeed, peak antibody levels were significantly correlated with later antibody levels (**figure S6E**). Memory B cell frequencies 1 week after the second dose were also correlated significantly with 3- and 6-month frequencies (**figure S6E**). Like antibodies and memory B cells, peak T cell responses after the second dose were predictive of later timepoints (**figure S6E**). Overall, these data suggest that the magnitude and trajectory of individual components of the immune response are patterned soon after the second vaccine dose in SARS-CoV-2 naïve individuals.

Finally, this dataset presented an opportunity to investigate the impact of “booster” mRNA vaccination in subjects with pre-existing immunity, in this case from a prior SARS-CoV-2 infection. To investigate the dynamics of boosted immunity, we examined the change in individual SARS-CoV-2-specific immune responses from pre-vaccine baseline levels. Booster vaccination modestly increased memory B cell and CD4+ T cell frequencies at 1 month, with a more robust increase in antibody levels (figure 6E). To investigate the contribution of pre-existing immune memory to these boosted antibody responses, we correlated the magnitude of pre-vaccine memory responses with the change in antibody levels after vaccination. The frequency of SARS-CoV-2-specific memory B cells was the only feature of pre-existing immunity that positively correlated with antibody responses after booster vaccination (figure 6F), consistent with a major role for memory B cells in recall responses. We next evaluated the decay kinetics of a boosted response. Boosting of Spike- and RBD-specific memory B cell and memory CD4+ T cell responses was transient and returned to pre-vaccination baseline by 3-6 months (figure 6E). CD8+ T cell responses were not boosted in SARS-CoV-2 immune subjects and decayed from peak at a comparable rate to SARS-CoV-2 naïve vaccinees (figure 6E). The boosting of anti-Spike and anti-RBD binding antibodies was also transient and returned to near baseline by 6 months (figure 6E). Only D614G and B.1.351 neutralizing antibody remained substantially above pre-vaccine baseline at 6 months (∼6.8 fold increase), but these titers were also declining over time. Notably, the decay rate of antibodies was similar between SARS-CoV-2 naïve and SARS-CoV-2 recovered vaccinees (figure 6E). Overall, these data suggest that boosting of pre-existing immunity with mRNA vaccination does not enhance already durable memory B cell or memory T cell responses. Rather, the benefit of booster vaccination in this setting may be limited to a significant but transient increase in antibody.

## CONCLUDING REMARKS

In summary, these studies demonstrate that mRNA vaccines induce durable immune memory to SARS-CoV-2 that continues to evolve over time. Examination of binding and neutralizing antibodies, memory B cells, and memory CD4+ and CD8+ T cells through 6 months post-vaccination highlighted several key features of mRNA vaccine-induced immunological memory. Specifically, these analyses revealed that SARS-CoV-2-specific memory B cell responses were robustly induced following mRNA vaccination and continue to increase in frequency for at least 6 months, even as circulating antibody levels declined in the same individuals. Moreover, mRNA vaccination generated highly mutated memory B cells that were capable of cross-binding VOCs, including B.1.1.7 (Alpha), B.1.351 (Beta), and B.1.617.2 (Delta). These vaccine-induced memory B cells were capable of rapidly generating new antibody responses and may have important contributions to protective immunity following exposure to SARS-CoV-2 in vaccinated subjects. mRNA vaccine-induced memory B cells also appeared to have a qualitative advantage at binding variants compared to memory B cells generated by mild COVID-19. Additionally, SARS-CoV-2-specific memory CD4+ T cells were relatively stable from 3-6 months post mRNA vaccination with a half-life of 194 days, and the vast majority of vaccinees had robust virus-specific CD4+ T cell responses at 6 months. Early CD4+ T cell responses correlated with 3- and 6-month humoral responses, highlighting a role for T cell immunity in shaping the overall response to vaccination. Together, these data identify durable cellular immunity for at least 6 months after mRNA vaccination with persistence of high-quality memory B cells and strong CD4+ T cell memory in most individuals.

These data may also provide context for understanding potential discrepancies in vaccine efficacy at preventing infection versus severe disease, hospitalization and death. Declining antibody titers over time likely reduce the potential for sterilizing or near sterilizing immunity. However, the durability of cellular immunity, here demonstrated for at least 6 months, may contribute to rapid recall responses that can limit initial viral replication and dissemination in the host, thereby preventing severe disease. Finally, by examining individuals with pre-existing immunity following natural infection, we were able to gain insights into the possible effects of booster vaccination. In this setting, boosting of pre-existing immunity with mRNA vaccination mainly resulted in a transient benefit to antibody titers with little-to-no long-term impact on cellular immune memory. Antibody decay rates were similar in SARS-CoV-2 naïve and recovered vaccinees, suggesting that additional vaccine doses will temporarily prolong antibody-mediated protection without fundamentally altering the underlying landscape of SARS-CoV-2 immune memory, which is characterized by durable memory B and CD4+ T cells even when antibody responses begin to wane. It will be important to examine whether similar dynamics exist following other types of immune boosting including homologous and heterologous vaccine boosting regimens. Nevertheless, these data provide evidence for durable immune memory at 6 months after mRNA vaccination and are relevant for interpreting epidemiological data on rates of infections in vaccinated populations and the implementation of booster vaccine strategies.

## LIMITATIONS OF THE STUDY

Despite the overall strengths of this study, including the large sample size and integrated measurement of multiple components of the antigen-specific adaptive immune response, there are several important limitations. First, the overall number of subjects, while substantial for studies with high depth of immune profiling, was still limited compared to epidemiological or phase 3 clinical trials. Second, it is possible that the timepoints sampled in this study do not perfectly capture the full kinetics of the response for each individual immune component. Additionally, the comparison of variant-specific immune memory induced by vaccination versus natural infection is limited to mild COVID-19 cases and does not include more severe disease. Timepoints for sampling of natural infection, although broadly consistent with the vaccination studies, were also not perfectly aligned with the date of actual infection as samples were longitudinally collected following a positive serology test rather than an acutely positive PCR test in most cases. Regarding CD8+ T cell responses, our AIM assay was effective at capturing peak responses after vaccination; however, this assay may not be sensitive enough to detect very low frequency CD8+ T cells at memory timepoints. Other approaches, such as tetramers, will be necessary in the future to further interrogate memory CD8+ T cell responses after vaccination. Finally, our cohort is overall skewed towards young healthy individuals. As such, the results described may not fully represent the durability of vaccine-induced immunity in older individuals or populations with chronic diseases and/or compromised immune systems, and future studies will be required to better quantify the immune response over time in these populations.

## METHODS

### Clinical Recruitment and Sample Collection

61 individuals (45 SARS-CoV-2 naïve, 16 SARS-CoV-2 recovered) were consented and enrolled in the study with approval from the University of Pennsylvania Institutional Review Board (IRB# 844642). All participants were otherwise healthy and based on self-reported health screening did not have any history of chronic health conditions. Subjects were stratified based on self-reported and laboratory evidence of a prior SARS-CoV-2 infection. All subjects received either Pfizer (BNT162b2) or Moderna (mRNA-1273) mRNA vaccines. Samples were collected at 6 timepoints: baseline, 2 weeks post-primary immunization, day of post-secondary immunization, 1 week post-secondary immunization, 3 months post-primary immunization, and 6 months post-primary immunization. 80-100mL of peripheral blood samples and clinical questionnaire data were collected at each study visit. A separate cohort of 19 SARS-CoV-2 convalescent individuals was used to compare vaccine-induced immune responses to immune responses upon natural SARS-CoV-2 infection. This cohort was a subset from a sero-monitoring study previously described (*34*) that was approved by the University of Pennsylvania Institutional Review Board (IRB# 842847). Recent or active SARS-CoV-2 infections were identified based on SARS-CoV-2 RBD antibody levels and/or SARS-COV-2 PCR testing. Longitudinal samples were collected from seropositive participants up to ∼200 days post seroconversion to study long-term immune responses. Full cohort and demographic information is provided in **table S1**.

### Peripheral Blood Sample Processing

Venous blood was collected into sodium heparin and EDTA tubes by standard phlebotomy. Blood tubes were centrifuged at 3000rpm for 15 minutes to separate plasma. Heparin and EDTA plasma were stored at −80°C for downstream antibody analysis. Remaining whole blood was diluted 1:1 with RPMI + 1% FBS + 2mM L-Glutamine + 100 U Penicillin/Streptomycin and layered onto SEPMATE tubes (STEMCELL Technologies) containing lymphoprep gradient (STEMCELL Technologies). SEPMATE tubes were centrifuged at 1200g for 10 minutes and the PBMC fraction was collected into new tubes. PBMCs were then washed with RPMI + 1% FBS + 2mM L-Glutamine + 100 U Penicillin/Streptomycin and treated with ACK lysis buffer (Thermo Fisher) for 5 minutes. Samples were washed again with RPMI + 1% FBS + 2mM L-Glutamine + 100 U Penicillin/Streptomycin, filtered with a 70µm filter, and counted using a Countess automated cell counter (Thermo Fisher). Aliquots containing 5-10×10^6^ PBMCs were cryopreserved in fresh 90% FBS 10% DMSO.

### Detection of SARS-CoV-2 Spike- and RBD-Specific Antibodies

Plasma samples were tested for SARS-CoV-2-specific antibody by enzyme-linked immunosorbent assay (ELISA) as described (*12, 48*). Plasmids encoding the recombinant full-length Spike protein and the RBD were provided by F. Krammer (Mt. Sinai) and purified by nickel-nitrilotriacetic acid resin (Qiagen). ELISA plates (Immulon 4 HBX, Thermo Fisher Scientific) were coated with PBS or 2 ug/mL recombinant protein and stored overnight at 4C. The next day, plates were washed with phosphate-buffered saline containing 0.1% Tween-20 (PBS-T) and blocked for 1 hour with PBS-T supplemented with 3% non-fat milk powder. Samples were heat-inactivated for 1 hour at 56C and diluted in PBS-T supplemented with 1% non-fat milk powder. After washing the plates with PBS-T, 50 uL diluted sample was added to each well. Plates were incubated for 2 hours and washed with PBS-T. Next, 50 uL of 1:5000 diluted goat anti-human IgG-HRP (Jackson ImmunoResearch Laboratories) or 1:1000 diluted goat anti-human IgM-HRP (SouthernBiotech) was added to each well and plates were incubated for 1 hour. Plates were washed with PBS-T before 50 uL SureBlue 3,3’,5,5’-tetramethylbenzidine substrate (KPL) was added to each well. After 5 minutes incubation, 25 uL of 250 mM hydrochloric acid was added to each well to stop the reaction. Plates were read with the SpectraMax 190 microplate reader (Molecular Devices) at an optical density (OD) of 450 nm. Monoclonal antibody CR3022 was included on each plate to convert OD values into relative antibody concentrations. Plasmids to express CR3022 were provided by I. Wilson (Scripps).

### Detection of SARS-CoV-2 Neutralizing Antibodies

293T cells were seeded for 24 hours at 5 × 10^6^ cells per 10 cm dish and were transfected using calcium phosphate with 35 μg of pCG1 SARS-CoV-2 S D614G delta18 or pCG1 SARS-CoV-2 S B.1.351 delta 18 expression plasmid encoding a codon optimized SARS-CoV-2 S gene with an 18-residue truncation in the cytoplasmic tail (kindly provided by Stefan Pohlmann). Variant sequences are provided below. 12 hours post transfection, cells were fed with fresh media containing 1mM sodium butyrate to increase expression of the transfected DNA. 24 hours after transfection, the SARS-CoV-2 Spike expressing cells were infected for 2 hours with VSV-G pseudotyped VSVΔG-RFP at an MOI of ∼1. Virus containing media was removed and the cells were re-fed with media without serum. Media containing the VSVΔG-RFP SARS-CoV-2 pseudotypes was harvested 28-30 hours after infection, clarified by centrifugation twice at 6000g then aliquoted, and stored at −80 °C until used for antibody neutralization analysis. All sera were heat-inactivated for 30 minutes at 55 ⁰C prior to use in the neutralization assay. Vero E6 cells stably expressing TMPRSS2 were seeded in 100 μl at 2.5×10^4^ cells/well in a 96 well collagen coated plate. The next day, 2-fold serially diluted serum samples were mixed with VSVΔG-RFP SARS-CoV-2 pseudotype virus (100-300 focus forming units/well) and incubated for 1 hour at 37⁰C. 1E9F9, a mouse anti-VSV Indiana G, was also included in this mixture at a concentration of 600 ng/ml (Absolute Antibody, Ab01402-2.0) to neutralize any potential VSV-G carryover virus. The serum-virus mixture was then used to replace the media on VeroE6 TMPRSS2 cells. 22 hours post-infection, the cells were washed and fixed with 4% paraformaldehyde before visualization on an S6 FluoroSpot Analyzer (CTL, Shaker Heights OH). Individual infected foci were enumerated and the values were compared to control wells without antibody. The focus reduction neutralization titer 50% (FRNT_50_) was measured as the greatest serum dilution at which focus count was reduced by at least 50% relative to control cells that were infected with pseudotype virus in the absence of human serum. FRNT_50_ titers for each sample were measured in at least two technical replicates and were reported for each sample as the geometric mean of the technical replicates.

| Variant | Mutations in Pseudovirus Construct |
| --- | --- |
| D614G (WT) | D614G |
| B.1.351 | L18F, D80A, D215G, R246I, K417N, E484K, N501Y, D614G, A701V |
| B.1.617.2 | T19R, G142D, del156-157, R158G, L452R, T478K, D614G, P681R, D950N |

### Detection and Phenotyping of SARS-CoV-2-Specific Memory B Cells

Antigen-specific B cells were detected using biotinylated proteins in combination with different streptavidin (SA)-fluorophore conjugates as described (*12*). All reagents are listed in **table S3**. Biotinylated proteins were multimerized with fluorescently labeled SA for 1 hour at 4C. Full-length Spike protein was mixed with SA-BV421 at a 10:1 mass ratio (e.g., 200ng Spike with 20ng SA; ∼4:1 molar ratio). Spike RBD was mixed with SA-APC at a 2:1 mass ratio (e.g., 25ng RBD with 12.5ng SA; ∼4:1 molar ratio). Biotinylated influenza HA pools were mixed with SA-PE at a 6.25:1 mass ratio (e.g., 100ng HA pool with 16ng SA; ∼6:1 molar ratio). Influenza HA antigens corresponding with the 2019 trivalent vaccine (A/Brisbane/02/2018/H1N1, B/Colorado/06/2017) were chosen as a historical antigen and were biotinylated using an EZ-Link Micro NHS-PEG4 Biotinylation Kit (Thermo Fisher) according to the manufacturer’s instructions. Excess biotin was subsequently removed using Zebra Spin Desalting Columns 7K MWCO (Thermo Fisher) and protein was quantified with a Pierce BCA Assay (Thermo Fisher). SA-BV711 was used as a decoy probe without biotinylated protein to gate out cells that non-specifically bind streptavidin. All experimental steps were performed in a 50/50 mixture of PBS + 2% FBS and Brilliant Buffer (BD Bioscience). Antigen probes for Spike, RBD, and HA were prepared individually and mixed together after multimerization with 5uM free D-biotin (Avidity LLC) to minimize potential cross-reactivity between probes. For staining, 5×10^6^ cryopreserved PBMC samples were prepared in a 96-well U-bottom plate. Cells were first stained with Fc block (Biolegend, 1:200) and Ghost 510 Viability Dye for 15 minutes at 4C. Cells were then washed and stained with 50uL antigen probe master mix containing 200ng Spike-BV421, 25ng RBD-APC, 100ng HA-PE, and 20ng SA-BV711 decoy for 1 hour at 4C. Following incubation with antigen probe, cells were washed again and stained with anti-CD3, anti-CD19, anti-CD20, anti-CD27, anti-CD38, anti-CD71, anti-IgD, anti-IgM, and anti-IgG for 30 minutes at 4C. After surface stain, cells were washed and fixed in 1% PFA overnight at 4C. Antigen-specific gates for B cell probe assays were set based on healthy donors stained without antigen probes (similar to an FMO control) and were kept the same for all experimental runs.

### Detection of Variant RBD, NTD, and S2-Specific Memory B Cells

Variant RBD, NTD, and S2-specific memory B cells were detected using a similar approach as described above. SARS-CoV-2 nucleocapsid was used as a vaccine-irrelevant antigen control. All reagents are listed in **table S3**. Probes were multimerized for 1.5 hours at the following ratios (all ∼4:1 molar ratios): 200ng full-length Spike protein was mixed with 20ng SA-BV421, 30ng N-terminal domain was mixed with 12ng SA-BV786, 25ng wild-type RBD was mixed with 12.5ng SA-BB515, 25ng B.1.1.7 RBD was mixed with 12.5ng SA-BV711, 25ng B.1.351 RBD was mixed with 12.5ng SA-PE, 25ng B.1.617.2 was mixed with 12.5ng SA-APC, 50ng S2 was mixed with 12ng SA-BUV737, 50ng nucleocapsid was mixed with 14ng SA-BV605. 12.5ng SA-BUV615 was used as a decoy probe. All antigen probes were multimerized separately and mixed together with 5uM free D-biotin. Prior to staining, total B cells were enriched from 20×10^6^ cryopreserved PBMC samples by negative selection using an EasySep human B cell isolation kit (STEMCELL, #17954). B cells were then prepared in a 96-well U-bottom plate and stained with Fc block and Ghost 510 Viability Dye as described above. Cells were washed and stained with 50uL antigen probe master mix for 1 hour at 4C. After probe staining, cells were washed again and stained with anti-CD3, anti-CD19, anti-CD27, anti-CD38, anti-IgD, and anti-IgG for 30 minutes at 4C. After surface stain, cells were washed and fixed in 1X Stabilizing Fixative (BD Biosciences) overnight at 4C.

For sorting, pre-enriched B cells were stained with Fc block and Ghost 510 Viability Dye, followed by full-length Spike, WT RBD, and B.1.351 RBD probes as described above. Cells were then stained for surface markers with anti-CD19, anti-CD20, anti-CD27, and anti-CD38, and anti-IgD. After surface stain, cells were washed and resuspended in PBS + 2% FBS for acquisition.

### In Vitro Differentiation of Memory B Cells to Antibody Secreting Cells

Memory B cells from bulk PBMC samples were differentiated into antibody secreting cells as described (*33*). Briefly, 1×10^6^ cryopreserved PBMCs were seeded in 1mL of complete RPMI media (RPMI + 10% FBS + 1% Pen/Strep) in 24-well plates. PBMCs were then stimulated with 1000U/mL recombinant human IL-2 and 2.5ug/mL R848 for 10 days. Supernatants were collected at the indicated timepoints, and anti-Spike IgG was quantified using a Human SARS-CoV-2 Spike (Trimer) IgG ELISA Kit (Invitrogen) according to the manufacturer’s instructions. RBD binding inhibition was measured using a SARS-CoV-2 Neutralizing Ab ELISA Kit (Invitrogen). For anti-Spike IgG experiments, culture supernatants were tested at 1:100 and 1:1000 dilutions. For RBD neutralization experiments, culture supernatants were tested without dilution and at a 1:2 dilution.

### Detection of SARS-CoV-2-Specific T Cells

SARS-CoV-2-specific T cells were detected using an activation induced marker assay. All reagents are listed in **table S4**. PBMCs were thawed by warming frozen cryovials in a 37°C water bath and resuspending cells in 10mL of RPMI supplemented with 10% FBS, 2mM L-Glutamine, 100 U/mL Penicillin, and 100 ug/mL Streptomycin (R10). Cells were washed once in R10, counted using a Countess automated cell counter (Thermo Fisher), and resuspended in fresh R10 to a density of 5×10^6^ cells/mL. For each condition, duplicate wells containing 1×10^6^ cells in 200uL were plated in 96-well round-bottom plates and rested overnight in a humidified incubator at 37°C, 5% CO2. After 16 hours, CD40 blocking antibody (0.5ug/mL final concentration) was added to cultures for 15 minutes prior to stimulation. Cells were then stimulated for 24 hours with costimulation (anti-human CD28/CD49d, BD Biosciences) and peptide megapools (CD4-S for all CD4+ T cell analyses, CD8-E for all CD8+ T cell analyses) at a final concentration of 1 ug/mL. Peptide megapools were prepared as previously described (*45, 46*). Matched unstimulated samples for each donor at each timepoint were treated with costimulation alone. 20 hours post-stimulation, antibodies targeting CXCR3, CCR7, CD40L, CD107a, CXCR5, and CCR6 were added to the culture along with monensin (GolgiStop, BD Biosciences) for a 4-hour stain at 37°C. After 4 hours, duplicate wells were pooled and cells were washed in PBS supplemented with 2% FBS (FACS buffer). Cells were stained for 10 minutes at room temperature with Ghost Dye Violet 510 and Fc receptor blocking solution (Human TruStain FcX™, BioLegend) and washed once in FACS buffer. Surface staining for 30 minutes at room temperature was then performed with antibodies directed against CD4, CD8, CD45RA, CD27, CD3, CD69, CD40L, CD200, OX40, and 41BB in FACS buffer. Cells were washed once in FACS buffer, fixed and permeabilized for 30 minutes at room temperature (eBioscience™ Foxp3 / Transcription Factor Fixation/Permeabilization Concentrate and Diluent), and washed once in 1X Permeabilization Buffer prior to staining for intracellular IFN-g overnight at 4°C. Cells were then washed again and resuspended in 1% paraformaldehyde in PBS prior to data acquisition.

All data from AIM expression assays were background-subtracted using paired unstimulated control samples. For memory T cell and helper T cell subsets, the AIM+ background frequency of non-naïve T cells was subtracted independently for each subset. AIM+ cells were identified from non-naïve T cell populations. AIM+ CD4+ T cells were defined by co-expression of CD200 and CD40L. AIM+ CD8+ T cells were defined by a Boolean analysis identifying cells expressing at least four of five markers: CD200, CD40L, 41BB, CD107a, and intracellular IFN-γ.

### Flow Cytometry and Cell Sorting

Samples were acquired on a BD Symphony A5 instrument. Standardized SPHERO rainbow beads (Spherotech) were used to track and adjust photomultiplier tubes over time. UltraComp eBeads (Thermo Fisher) were used for compensation. Up to 5×10^6^ cells were acquired per sample. Data were analyzed using FlowJo v10 (BD Bioscience).

### B Cell Receptor Sequencing

#### Library Preparation

DNA was extracted from sorted cells using a Gentra Puregene Cell kit (Qiagen, catalog no. 158767). Immunoglobulin heavy-chain family–specific PCRs were performed on genomic DNA samples using primers in FR1 and JH as described previously (*41, 49*). Two biological replicates were run on all samples. Sequencing was performed in the Human Immunology Core Facility at the University of Pennsylvania using an Illumina 2× 300-bp paired-end kit (Illumina MiSeq Reagent Kit v3, 600-cycle, Illumina MS-102-3003).

#### IGH Sequence Analysis

Reads from the Illumina MiSeq were filtered, annotated, and grouped into clones as described previously (*12, 50*). Briefly, pRESTO v0.6.0 (*51*) was used to align paired end reads, remove short and low-quality reads, and mask low-quality bases with *N*s to avoid skewing SHM and lineage analyses. Sequences which passed this process were aligned and annotated with IgBLAST v1.17.0 (*52*). The annotated sequences were then imported into ImmuneDB v0.29.10 (*53, 54*) for clonal inference, lineage construction, and downstream processing. For clonal inference, sequences with the same VH, VJ, and CDR3 length from each donor were hierarchically clustered. Those sequences sharing 85% or higher similarity in their CDR3 amino-acid sequence were subsequently grouped into clones.

#### Lineage Construction & Visualization

A lineage for each clone was constructed with ImmuneDB, a lineage was constructed as described in (*54*). ete3 (*55*) was used to visualize the lineages where each node represents a unique sequence, the size of a node represents its relative copy number fraction in the clone, the integer next to each node represents the number of mutations from the preceding vertical node.

#### Overlap and Overlapping Clone SHM Analysis

Clones were filtered based on size using a copy number cut-off of 50% of the mean copy number frequency per clone (≥50% mcf) in each sequencing library. Large clones were analyzed for overlap between the sorted subpopulations and numbers of clones in each sorted population were visualized using Venn Diagrams. SHM within ≥50% mcf clones that overlapped between WT RBD and cross-binders (RBD++) was analyzed within each sequence in the clone and averaged for all of the sequence copies that were found in each clone. SHM data were aggregated across all subjects with each clone counting once irrespective of its size. Clonal SHM levels were visualized as categorical variables (pie chart) and as frequencies (histogram plot).

#### Data Availability

Raw sequencing data for all donors and subsets is available on SRA under BioProject PRJNA752617.

### High Dimensional Analysis and Statistics

All data were analyzed using custom scripts in R and visualized using RStudio. For heatmaps, all data were scaled by column (z-score normalization). For construction of UMAPs, antibody and cell frequency data were log10 transformed and scaled by column (z-score normalization) prior to generating UMAP coordinates. Statistical tests are indicated in the corresponding figure legends. All tests were performed two-sided with a nominal significance threshold of p < 0.05. Holm correction was performed in all cases of multiple comparisons. Unpaired tests were used for comparisons between timepoints unless otherwise indicated as some participants were missing samples from individual timepoints. * indicates p < 0.05, ** indicates p < 0.01, *** indicates p < 0.001, **** indicates p < 0.0001. Source code and data files are available upon request from the authors.

## ACKNOWLEDGEMENTS

We thank the study participants for their generosity in making the study possible. We also thank Shane Crotty and members of the Wherry lab for helpful discussions and feedback, as well as the Flow Cytometry Core at the University of Pennsylvania for technical support. This work was supported by grants from the NIH AI105343, AI082630, AI108545, AI155577, AI149680 (to EJW), AI152236 (to PB), HL143613 (to JRG), P30-AI0450080 (to ELP), T32 AR076951-01 (to SAA), T32 CA009140 (to JRG and DM), T32 AI055400 (to PH), U19AI082630 (to SEH and EJW), funding from the Allen Institute for Immunology (to SAA, EJW), Cancer Research Institute-Mark Foundation Fellowship (to JRG), Chen Family Research Fund (to SAA), the Parker Institute for Cancer Immunotherapy (to JRG, EJW), the Penn Center for Research on Coronavirus and Other Emerging Pathogens (to PB), the University of Pennsylvania Perelman School of Medicine COVID Fund (to RRG, EJW), the University of Pennsylvania Perelman School of Medicine 21^st^ Century Scholar Fund (to RRG), and a philanthropic gift from Jeffrey Lurie, Joel Embiid, Josh Harris, and David Blitzer (to SEH). Work in the Wherry lab is supported by the Parker Institute for Cancer Immunotherapy. This work was also supported by NIH contract Nr. 75N9301900065 (to DW, AS).

## AUTHOR CONTRIBUTIONS

RRG, MMP, and EJW designed the study. RRG, MMP, SAA, DM, WM, KL, SG, LKC, PH, SD, MEW, CMM, MA, NT, and EMD carried out experiments. RRG, SAA, JD, SL, and OK were involved in clinical recruitment and sample collection. WM, AMR, AR, DSK, DAO, JRG, MPD, and ELP provided expertise on statistical analyses. RRG, MMP, DM, and AEB contributed to the methodology. RRG, MMP, AP, AH, HS, SH, SK, JTH, JCW, and SA processed peripheral blood samples and managed the sample database. IF, AG, DW, and AS provided key samples and/or reagents. ELP, ARG and EJW supervised the study. All authors participated in data analysis and interpretation. RRG, MMP, ELP, and EJW wrote the manuscript.

The UPenn COVID Processing Unit included individuals from diverse laboratories at the University of Pennsylvania who volunteered their time and effort to enable study of COVID-19 patients during the pandemic: Sharon Adamski, Zahidul Alam, Mary M. Addison, Katelyn T. Byrne, Aditi Chandra, Hélène C. Descamps, Nicholas Han, Yaroslav Kaminskiy, Shane C. Kammerman, Justin Kim, Allison R. Greenplate, Jacob T. Hamilton, Nune Markosyan, Julia Han Noll, Dalia K. Omran, Ajinkya Pattekar, Eric Perkey, Elizabeth M. Prager, Dana Pueschl, Austin Rennels, Jennifer B. Shah, Jake S. Shilan, Nils Wilhausen, Ashley N. Vanderbeck. All are affiliated with the University of Pennsylvania Perelman School of Medicine.

