## Supplemental Information for "mRNA Vaccination Induces Durable Immune Memory to SARS-CoV-2 with Continued Evolution to Variants of Concern"

**Figure S1. Gating strategy for SARS-CoV-2-specific memory B cells.** For **panel 1**, lymphocytes were first identified based on forward- and side-scatter from bulk PBMC samples. Singlets were excluded by FSC-A/FSC-H and FSC-A/FSC-W. Dead cells were excluded using Ghost 510 viability dye. Total B cells were then identified as CD3- CD19+ cells. Naïve B cells were identified as IgD+ CD27- B cells and excluded from downstream analysis. Memory B cells were subsequently identified from non-naïve B cells as CD20+ CD38lo/int cells. A BV711 decoy probe was used to gate out memory B cells that non-specifically bound streptavidin. Spike- and HA-binding were then quantified on decoy- memory B cells. Binding to RBD probe was also measured on Spike+ memory B cells. IgG, IgM, and IgA isotypes were evaluated for both Spike+ and Spike+ RBD+ memory B cells. CD71 was measured as an activation marker on Spike+ memory B cells. For **panel 2**, total B cells were enriched by negative selection from PBMC samples prior to staining. Live, non-naïve B cells were identified as described above. Plasmablasts were then identified as CD27+ CD38+ non-naïve B cells and were excluded from downstream analysis. Decoy- cells were excluded as described above. Spike- and nucleocapsid-specific B cells were identified based on binding to corresponding probes. Spike+ memory cells were then analyzed for co-binding to N-terminal domain (NTD) or S2 domain probes. Memory B cells that were Spike+ but NTD- and S2- were subsequently analyzed for co-binding to a panel of variant RBD probes, including wild-type (WT), B.1.1.7, B.1.351, and B.1.617.2 RBDs. IgG expression was evaluated for all antigen-specific populations.

**Figure S2. Class-switching of Spike+ and Spike+ RBD+ memory B cells after mRNA vaccination. A)** Percent IgG+ of Spike+ and **B)** Spike+ RBD+ memory B cells over time after mRNA vaccination. **C)** Summary statistics for % IgG+, % IgM+, and % other isotype+ of SARS-CoV-2-specific memory B cells over time. Outer rings represent total Spike+ memory B cells, inner rings represent Spike+ RBD+ memory B cells.

**Figure S3. Extended analysis of SARS-CoV-2-specific memory B cell frequencies and class-switching after mRNA vaccination. A)** Frequency of SARS-CoV-2 antigen-specific memory B cells over time after mRNA vaccination or infection. Data are represented as a percentage of total B cells. **B)** Cross sectional analysis of antigen-specific memory B cell frequencies at 6 months post-vaccination or sero-positivity. Pre-immune baseline samples are also shown as a control. **C)** Class-switching of RBD-binding cells over time after mRNA vaccination or infection. Data are represented as the percent of cells that are IgG+. **D)** Cross sectional analysis of class-switching to IgG at 6 months post-vaccination or sero-positivity.

**Figure S4. Extended analysis of variant-binding BCR sequencing data. A)** Individual flow plots of sorted memory B cell populations in 8 SARS-CoV-2 naïve and 4 SARS-CoV-2 recovered individuals. **B)** Histogram of VH gene usage across different antigen binding populations. Data are represented as a percentage of the overall clones for a given antigen-binding population. **C)** Individual SHM distributions of memory B cell clones for SARS-CoV-2 naïve and recovered individuals. Data are represented as the percent of mutated VH gene nucleotides. **D)** Venn diagram of clonal lineages that are shared between RBD-, WT RBD+ and RBD++ populations. Data were filtered based on larger clones with ≥ 50% mean copy number frequency in each sequencing library.

**Figure S5. Gating strategy for SARS-CoV-2-specific memory T cells.** Lymphocytes were first identified based on forward- and side-scatter from bulk PBMC samples. Singlets were excluded by FSC-A/FSC-H and SSC-A/SSC-H. Total T cells were identified as Live/Dead- CD3+. CD4 and CD8 T cells were then identified from total T cells. For both CD4+ and CD8+ T cells, naïve cells were identified as CD45RA+ CD27+ and excluded from downstream analysis. Memory subsets were defined based on a combination of CD45A, CD27, and CCR7 expression. CD4+ helper subsets were defined based on CCR6, CXCR3, and CXCR5 chemokine receptor expression. AIM+ CD4+ T cells were identified based on co-expression of CD40L and CD200. AIM+ CD8+ T cells were identified based on co-expression of at least 4 of 5 activation induced markers (intracellular IFN-$\gamma$, 41BB, CD40L, CD107a, CD200).

**Figure S6. Immune correlations after mRNA vaccination. A)** Relationship between age and overall vaccine response in SARS-CoV-2 naïve subjects. 6 month post-vaccination samples are colored by age and projected onto the UMAP coordinates from figure 6. **B)** Correlation between individual immune parameters and age at 6 months post-vaccination in SARS-CoV-2 naïve subjects. **C)** Relationship between sex and overall vaccine response in SARS-CoV-2 naïve subjects. 6 month post-vaccination samples are colored by sex and projected onto the UMAP coordinates from figure 6. **D)** Correlation between T cell responses and variant-specific humoral responses over time in SARS-CoV-2 naïve subjects. **E)** Correlation between peak antibody, memory B, and memory T cell responses 1 week after the second vaccine dose with later responses at 3 and 6 months post-vaccination in SARS-CoV-2 naïve subjects. All statistics were calculated using non-parametric Spearman rank correlation.

| | | SARS-CoV-2 Naïve | SARS-CoV-2 Recovered | Natural Infection (HCW) |
|---|---|---|---|---|
| **Total** | Number | 45 | 16 | 19 |
| **Age** | Average (Years) | 36.9 | 38.3 | 35.2 |
| | 20-30 | 15 | 4 | 8 |
| | 30-40 | 14 | 6 | 7 |
| | 40-50 | 9 | 2 | 2 |
| | 50+ | 7 | 4 | 2 |
| **Sex** | Male | 21 | 10 | 5 |
| | Female | 24 | 6 | 14 |
| **Race/Ethnicity** | White - Non-Hispanic/Latino | 27 | 7 | 17 |
| | White - Hispanic/Latino | 4 | 1 | 0 |
| | Asian | 8 | 6 | 1 |
| | Black | 2 | 1 | 1 |
| | Native | 0 | 1 | 0 |
| | Other | 1 | 0 | 0 |
| **Vaccine Type** | Pfizer | 42 | 12 | -- |
| | Moderna | 3 | 4 | -- |

**Table S1. Demographic Information for University of Pennsylvania Healthy COVID Vaccine and Healthcare Worker (HCW) Sero-Monitoring Studies.** Number of participants, age, sex, race/ethnicity, and vaccine type are indicated.

| Sample ID | Recovered | Population | Total DNA (ng) | Input DNA (ng) | Copies | Unique | In Frame | # Clones | Average VH Identity | CDR3 Length (NT) |
|---|---|---|---|---|---|---|---|---|---|---|
| N1 | No | Naïve B | 4160 | 200 | 140801 | 113799 | 0.841876 | 14242 | 0.984176 | 55.05961 |
| N1 | No | Spike+ RBD- | 38.64 | 35.88 | 164565 | 131819 | 0.852932 | 6684 | 0.959396 | 52.42774 |
| N1 | No | Spike+ WT RBD+ | 10.248 | 9.516 | 237325 | 121844 | 0.829343 | 2297 | 0.96788 | 50.98389 |
| N1 | No | Spike+ RBD++ | 3.892 | 3.614 | 212006 | 106777 | 0.808652 | 1803 | 0.957687 | 50.4376 |
| N2 | No | Naïve B | 3848 | 200 | 160856 | 136888 | 0.82964 | 18772 | 0.981181 | 56.12348 |
| N2 | No | Spike+ RBD- | 50.4 | 46.8 | 138341 | 115801 | 0.86341 | 7043 | 0.949157 | 55.41389 |
| N2 | No | Spike+ WT RBD+ | 22.82 | 21.19 | 185765 | 121679 | 0.827671 | 4474 | 0.969472 | 55.88869 |
| N2 | No | Spike+ RBD++ | 6.188 | 5.746 | 193081 | 100240 | 0.83583 | 1669 | 0.954832 | 53.36968 |
| N3 | No | Naïve B | 4108 | 200 | 166692 | 133169 | 0.836719 | 15507 | 0.982458 | 55.35513 |
| N3 | No | Spike+ RBD- | 22.26 | 20.67 | 159970 | 104626 | 0.855542 | 5891 | 0.956951 | 52.64828 |
| N3 | No | Spike+ WT RBD+ | 32.2 | 29.9 | 172876 | 127480 | 0.849711 | 6401 | 0.966868 | 54.50133 |
| N3 | No | Spike+ RBD++ | 3.976 | 3.692 | 176383 | 88649 | 0.823144 | 1374 | 0.961225 | 51.42868 |
| N4 | No | Naïve B | 3406 | 200 | 201283 | 153755 | 0.822365 | 15363 | 0.986036 | 57.14008 |
| N4 | No | Spike+ RBD- | 50.4 | 46.8 | 205448 | 149192 | 0.823989 | 5119 | 0.969826 | 54.63098 |
| N4 | No | Spike+ WT RBD+ | | | 195133 | 34123 | 0.700326 | 307 | 0.970296 | 54.71987 |
| N4 | No | Spike+ RBD++ | 4.256 | 3.952 | 235928 | 95887 | 0.796986 | 1128 | 0.960086 | 51.63564 |
| N5 | No | Naïve B | 532 | 200 | 151633 | 117639 | 0.853735 | 20456 | 0.962536 | 53.97961 |
| N5 | No | Spike+ RBD- | | | 11048 | 2265 | 0.571429 | 21 | 0.954252 | 59.66667 |
| N5 | No | Spike+ WT RBD+ | 5.768 | 5.356 | 130442 | 52200 | 0.812448 | 1205 | 0.965277 | 54.18755 |
| N5 | No | Spike+ RBD++ | | | 148307 | 21122 | 0.681373 | 204 | 0.974878 | 51.85294 |
| N6 | No | Naïve B | 535.5 | 200 | 206026 | 166095 | 0.848853 | 19180 | 0.97315 | 54.13264 |
| N6 | No | Spike+ RBD- | 5.292 | 4.914 | 180711 | 85411 | 0.819632 | 1630 | 0.975975 | 53.02393 |
| N6 | No | Spike+ WT RBD+ | 4.984 | 4.628 | 243434 | 85176 | 0.814755 | 1247 | 0.97978 | 51.36889 |
| N6 | No | Spike+ RBD++ | | | 145249 | 37847 | 0.759709 | 412 | 0.979334 | 52.58981 |
| N7 | No | Naïve B | 483 | 200 | 222335 | 177357 | 0.852768 | 22040 | 0.971328 | 53.32114 |
| N7 | No | Spike+ RBD- | 2.576 | 2.392 | 277624 | 98253 | 0.789958 | 1195 | 0.976346 | 50.7046 |
| N7 | No | Spike+ WT RBD+ | 8.092 | 7.514 | 286754 | 140707 | 0.801635 | 2324 | 0.974052 | 52.3352 |
| N7 | No | Spike+ RBD++ | 1.764 | 1.638 | 303988 | 71141 | 0.773611 | 720 | 0.979057 | 49.67778 |
| N8 | No | Naïve B | 343 | 127.4 | 133517 | 103687 | 0.861062 | 13632 | 0.972975 | 53.99831 |
| N8 | No | Spike+ RBD- | | | 195913 | 44220 | 0.665782 | 377 | 0.97288 | 55.687 |
| N8 | No | Spike+ WT RBD+ | | | 131695 | 28512 | 0.8 | 275 | 0.969267 | 50.82182 |
| N8 | No | Spike+ RBD++ | | | 19648 | 3645 | 0.815789 | 76 | 0.970741 | 45.21053 |
| R1 | Yes | Naïve B | 5564 | 200 | 166874 | 129000 | 0.837699 | 24054 | 0.985936 | 55.67278 |
| R1 | Yes | Spike+ RBD- | 24.5 | 22.75 | 184946 | 100096 | 0.851991 | 3466 | 0.945325 | 53.27986 |
| R1 | Yes | Spike+ WT RBD+ | 25.76 | 23.92 | 177969 | 107237 | 0.826538 | 5494 | 0.979587 | 56.57117 |
| R1 | Yes | Spike+ RBD++ | | | 191869 | 38316 | 0.699115 | 339 | 0.982212 | 54.56637 |
| R2 | Yes | Naïve B | 3666 | 200 | 150269 | 135921 | 0.861352 | 46535 | 0.985534 | 53.59173 |
| R2 | Yes | Spike+ RBD- | 21.14 | 19.63 | 156519 | 99743 | 0.864045 | 4222 | 0.955645 | 53.29725 |
| R2 | Yes | Spike+ WT RBD+ | 3.192 | 2.964 | 249006 | 86313 | 0.811163 | 1075 | 0.963926 | 54.00837 |
| R2 | Yes | Spike+ RBD++ | | | 254772 | 64204 | 0.843662 | 710 | 0.95905 | 53.49437 |
| R3 | Yes | Naïve B | 5239 | 200 | 243860 | 191104 | 0.841811 | 31671 | 0.986992 | 56.29873 |
| R3 | Yes | Spike+ RBD- | 24.22 | 22.49 | 133725 | 83873 | 0.863046 | 3848 | 0.950538 | 51.55249 |
| R3 | Yes | Spike+ WT RBD+ | 7.588 | 7.046 | 272563 | 120491 | 0.818267 | 2124 | 0.970827 | 55.60734 |
| R3 | Yes | Spike+ RBD++ | | | 245955 | 60802 | 0.709677 | 589 | 0.987109 | 53.60272 |
| R4 | Yes | Naïve B | 4186 | 200 | 193799 | 155701 | 0.83858 | 22680 | 0.981337 | 55.67928 |
| R4 | Yes | Spike+ RBD- | 24.78 | 23.01 | 195890 | 119951 | 0.849797 | 3948 | 0.955369 | 53.78343 |
| R4 | Yes | Spike+ WT RBD+ | 14.7 | 13.65 | 209686 | 105495 | 0.846287 | 3474 | 0.967402 | 54.31031 |
| R4 | Yes | Spike+ RBD++ | 5.936 | 5.512 | 160571 | 59064 | 0.851287 | 1049 | 0.95557 | 52.45281 |

**Table S2. BCR Sequencing Metadata.** B cell receptor sequencing metadata. Sample ID indicates subject; Recovered indicates prior COVID-19. Two independent PCR amplifications (biological replicates) were performed for each sample; Input DNA per replicate; Number of valid sequence copies (passing length and other QC filters, see methods); Clones are defined as sequences that share the same VH, JH, CDR3 length and are at least 85% identical in the third complementarity determining region (CDR3) amino acid sequence; Clones with only 1 copy at the subject level are excluded; Average VH identity compared to the nearest germline VH gene (average identity was calculated for each clone and then averaged across clones with each clone counted once per sample); CDR3 length in nucleotides (nt). Productive rearrangements only.

| Reagent | Vendor | Identifier | Concentration |
|---|---|---|---|
| **Panel 1 - B Cell Probe** | | | |
| SARS-CoV-2 Biotinylated Full Length Spike | R&D Systems | BT10549-050 | 200ng |
| SARS-CoV-2 Biotinylated Full Length Spike | R&D Systems | BT10500-050 | 25ng |
| HA(ΔTM)(A/Brisbane/02/2018)(H1N1) | Immune Tech | IT-003-00110ΔTMp | 50ng |
| HA(ΔTM)(B/Colorado/06/2017) | Immune Tech | IT-003-B21ΔTMp | 50ng |
| BV421 Streptavidin | Biolegend | 405226 | 20ng |
| BV711 Streptavidin | BD Biosciences | 563262 | 20ng |
| PE Streptavidin | Biolegend | 405203 | 16ng |
| APC Streptavidin | Biolegend | 405207 | 12.5ng |
| Ghost Viability Dye Violet 510 | Tonbo | 13-0870-T100 | 1:600 |
| BUV563 anti-CD3 | BD Biosciences | 748569 | 1:200 |
| BV750 anti-CD19 | Biolegend | 302262 | 1:100 |
| BUV805 anti-CD20 | BD Biosciences | 612905 | 1:500 |
| BUV395 anti-CD27 | BD Biosciences | 563815 | 1:200 |
| BUV661 anti-CD38 | BD Biosciences | 612969 | 1:200 |
| APC-H7 anti-CD71 | BD Biosciences | 563671 | 1:50 |
| FITC anti-IgA | Miltenyi | 130-113-475 | 1:400 |
| BV480 anti-IgD | BD Biosciences | 566138 | 1:50 |
| PE-Cy7 anti-IgG | Biolegend | 410722 | 1:400 |
| PerCP/Cy5.5 anti-IgM | Biolegend | 314512 | 1:400 |
| **Panel 2 - Variant B Cell Probe** | | | |
| SARS-CoV-2 Biotinylated Full Length Spike | R&D Systems | AVI10549-050 | 200ng |
| SARS-CoV-2 Biotinylated RBD | Acro Biosystems | SPD-C82E9-25ug | 25ng |
| SARS-CoV-2 Biotinylated RBD (N501Y) | Acro Biosystems | SPD-C82E6-25ug | 25ng |
| SARS-CoV-2 Biotinylated RBD (K417N/E484K/N501Y) | Acro Biosystems | SPD-C82E5-25ug | 25ng |
| SARS-CoV-2 Biotinylated RBD (L452R/K478N) | Acro Biosystems | SPD-C82Ed-25ug | 25ng |
| SARS-CoV-2 Biotinylated N-Terminal Domain | Sino Biological | 40591-V49H-B | 30ng |
| SARS-CoV-2 Biotinylated S2 | Acro Biosystems | S2N-C52E8-25ug | 50ng |
| SARS-CoV-2 Biotinylated Nucleocapsid | R&D Systems | BT10474-050 | 50ng |
| BV421 Streptavidin | Biolegend | 405226 | 20ng |
| BV605 Streptavidin | Biolegend | 405229 | 14ng |
| BV711 Streptavidin | BD Biosciences | 563262 | 12.5ng |
| BV786 Streptavidin | BD Biosciences | 563858 | 12ng |
| BUV615 Streptavidin | BD Biosciences | 613013 | 12.5ng |
| BUV737 Streptavidin | BD Biosciences | 612775 | 12ng |
| BB515 Streptavidin | BD Biosciences | 564453 | 12.5ng |
| PE Streptavidin | Biolegend | 405203 | 12.5ng |
| APC Streptavidin | Biolegend | 405207 | 12.5ng |
| Ghost Viability Dye Violet 510 | Tonbo | 13-0870-T100 | 1:600 |
| BUV563 anti-CD3 | BD Biosciences | 748569 | 1:200 |
| BV750 anti-CD19 | Biolegend | 302262 | 1:100 |
| BUV395 anti-CD27 | BD Biosciences | 563815 | 1:200 |
| BUV661 anti-CD38 | BD Biosciences | 612969 | 1:200 |
| BV480 anti-IgD | BD Biosciences | 566138 | 1:50 |
| APC-H7 anti-IgG | BD Biosciences | 561297 | 1:100 |
| **Panel 3 - Variant-Specific B Cell Sorting** | | | |
| SARS-CoV-2 Biotinylated Full Length Spike | R&D Systems | BT10549-050 | 200ng |
| SARS-CoV-2 Biotinylated RBD | Sino Biological | 40592-V08B-B | 25ng |

| | | | |
|---|---|---|---|
| SARS-CoV-2 Biotinylated RBD (K417N/E484K/N501Y) | Sino Biological | 40592-V08H85-B | 25ng |
| BV421 Streptavidin | Biolegend | 405226 | 20ng |
| AF488 Streptavidin | Biolegend | 405235 | 20ng |
| PE Streptavidin | Biolegend | 405203 | 12.5ng |
| APC Streptavidin | Biolegend | 405207 | 12.5ng |
| Ghost Viability Dye Violet 510 | Tonbo | 13-0870-T100 | 1:600 |
| APC-Cy7 anti-CD19 | BD Biosciences | 557791 | 1:200 |
| BV650 anti-CD20 | Biolegend | 302336 | 1:200 |
| BV785 anti-CD27 | Biolegend | 302832 | 1:66 |
| PE-Cy7 anti-CD38 | eBioscience | 25-0389-42 | 1:200 |
| PE-CF594 anti-IgD | BD Biosciences | 562540 | 1:50 |

**Table S3. Reagents for Memory B Cell Analysis.** Reagent, vendor, catalog number, and concentration/dilution are indicated.

| Reagent | Vendor | Identifier |
|---|---|---|
| **Flow Cytometry Antibodies** | | |
| BUV395 CD4 | BD Biosciences | Cat#563550 |
| BUV496 CD8 | BD Biosciences | Cat#612943 |
| BUV615 CD45RA | BD Biosciences | Cat#751555 |
| BUV737 CD27 | BD Biosciences | Cat#612829 |
| BUV805 CD3 | BD Biosciences | Cat#612896 |
| BV421 CXCR3 | Biolegend | Cat#353716 |
| BV650 CCR7 | Biolegend | Cat#353234 |
| BV605 CD69 | Biolegend | Cat#310938 |
| BV711 CD40L | Biolegend | Cat#310838 |
| BV785 CD107a | Biolegend | Cat#328644 |
| FITC IFNy | Biolegend | Cat#502515 |
| PE CD200 | Biolegend | Cat#399804 |
| PE-Cy7 OX40 | Biolegend | Cat#350012 |
| AF647 41BB | Biolegend | Cat#309810 |
| APC-R700 CXCR5 | BD Biosciences | Cat#565191 |
| APC-Cy7 CCR6 | Biolegend | Cat#353432 |
| **Peptides** | | |
| CD4-S peptide Megapool | Synthetic Biomolecules (aka A&A) | http://www.syntheticbiomolecules.com/ |
| CD8-E peptide Megapool | Synthetic Biomolecules (aka A&A) | http://www.syntheticbiomolecules.com/ |
| **Other** | | |
| Ghost Dye Violet 510 | Tonbo | Cat#13-0870-T500 |
| GolgiStop (Containing Monensin) | BD Biosciences | Cat#51-2092K7 |
| CD40 Antibody, anti-human, pure-functional grade | Miltenyi Biotech | Cat#130-094-133 |
| Anti-Human CD28/CD49d Purified | BD Biosciences | Cat#347690 |
| Human TruStain FcX™ (Fc Receptor Blocking Solution) | Biolegend | Cat#422302 |
| Foxp3 / Transcription Factor Fixation/Permeabilization Concentrate and Diluent | eBioscience | Cat#00-5521-00 |

**Table S4. Reagents for Memory T Cell Analysis.** Reagent, vendor, and catalog number are indicated.

### Figure S1

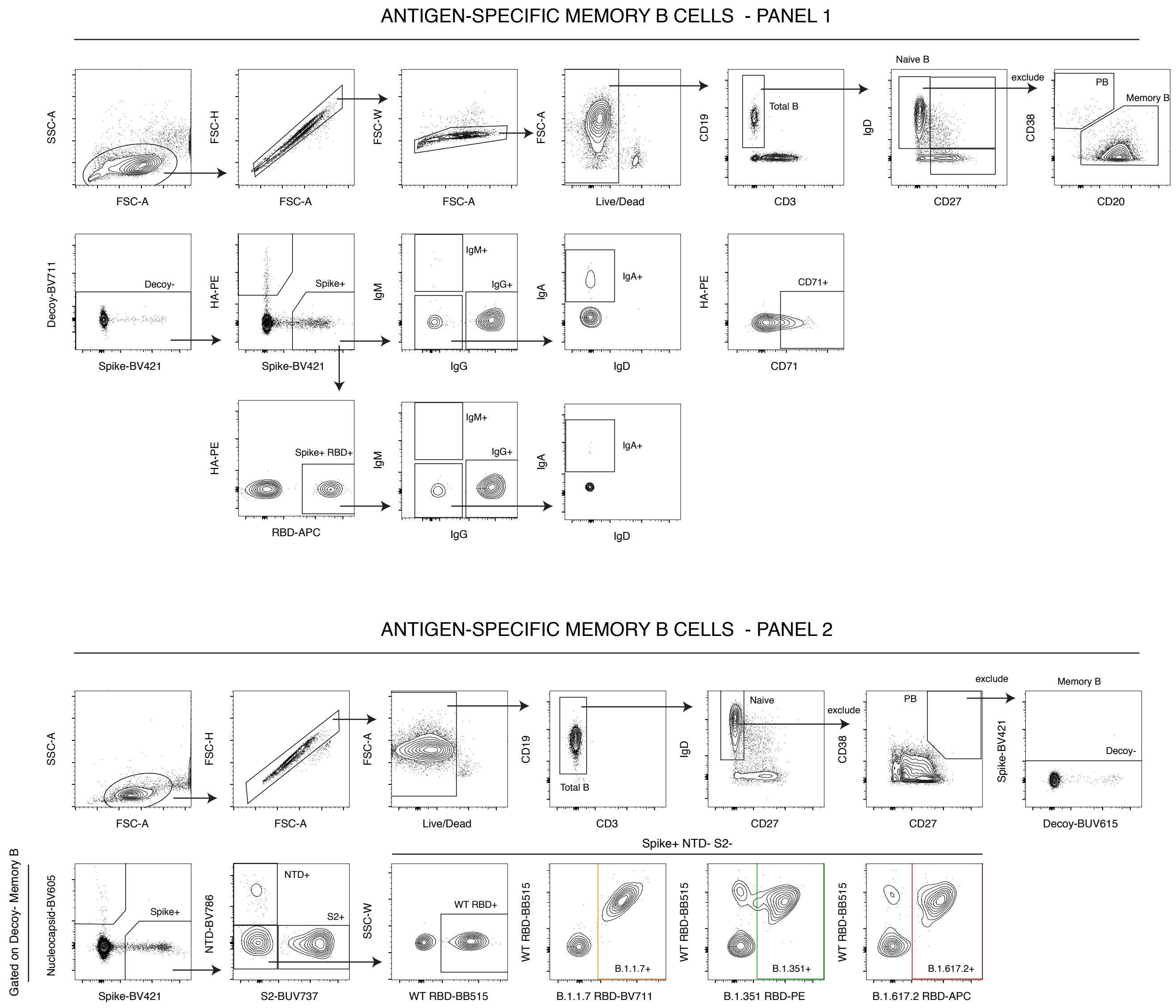
ANTIGEN-SPECIFIC MEMORY B CELLS - PANEL 1
SSC-A
FSC-A
FSC-H
FSC-W
Live/Dead
CD19
CD3
Total B
Naive B
IgD
CD27
exclude
PB
Memory B
CD38
CD20
Decoy-BV711
Decoy-
Spike-BV421
HA-PE
Spike+
IgM
IgM+
IgG+
IgG
IgA
IgA+
CD71+
CD71
Spike+ RBD+
RBD-APC
ANTIGEN-SPECIFIC MEMORY B CELLS - PANEL 2
Naive
Decoy-
Decoy-BUV615
Spike-BV421
Gated on Decoy- Memory B
Nucleocapsid-BV605
NTD-BV786
NTD+
S2+
S2-BUV737
Spike+ NTD- S2-
SSC-W
WT RBD+
WT RBD-BB515
B.1.1.7+
B.1.1.7 RBD-BV711
B.1.351+
B.1.351 RBD-PE
B.1.617.2+
B.1.617.2 RBD-APC

### Figure S2

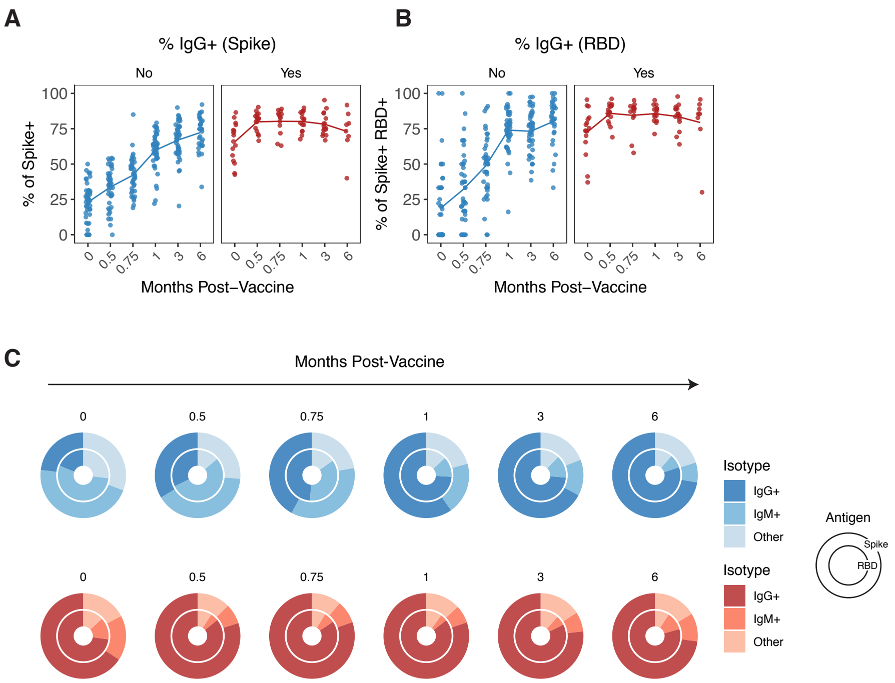
A
% IgG+ (Spike)
No
Yes
% of Spike+
100
75
50
25
0
0
0.5
0.75
1
3
6
Months Post–Vaccine
B
% IgG+ (RBD)
No
Yes
% of Spike+ RBD+
Months Post–Vaccine
C
Months Post-Vaccine
0
0.5
0.75
1
3
6
Isotype
IgG+
IgM+
Other
Antigen
Spike
RBD

### Figure S3

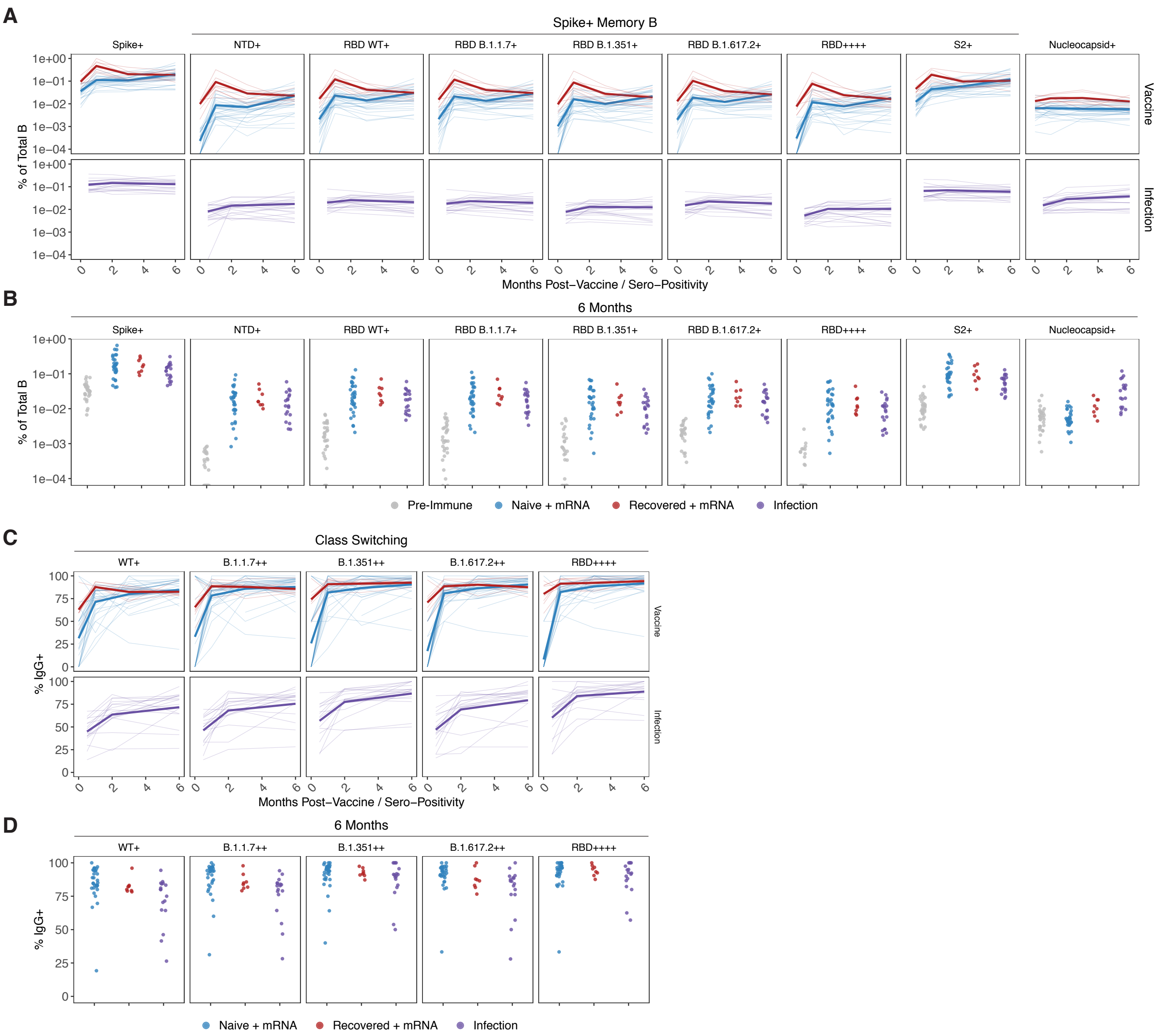
A
Spike+ Memory B
Spike+
NTD+
RBD WT+
RBD B.1.1.7+
RBD B.1.351+
RBD B.1.617.2+
RBD++++
S2+
Nucleocapsid+
Vaccine
Infection
% of Total B
Months Post−Vaccine / Sero−Positivity
B
6 Months
Pre-Immune
Naive + mRNA
Recovered + mRNA
Infection
C
Class Switching
WT+
B.1.1.7++
B.1.351++
B.1.617.2++
RBD++++
% IgG+
D
6 Months

### Figure S4

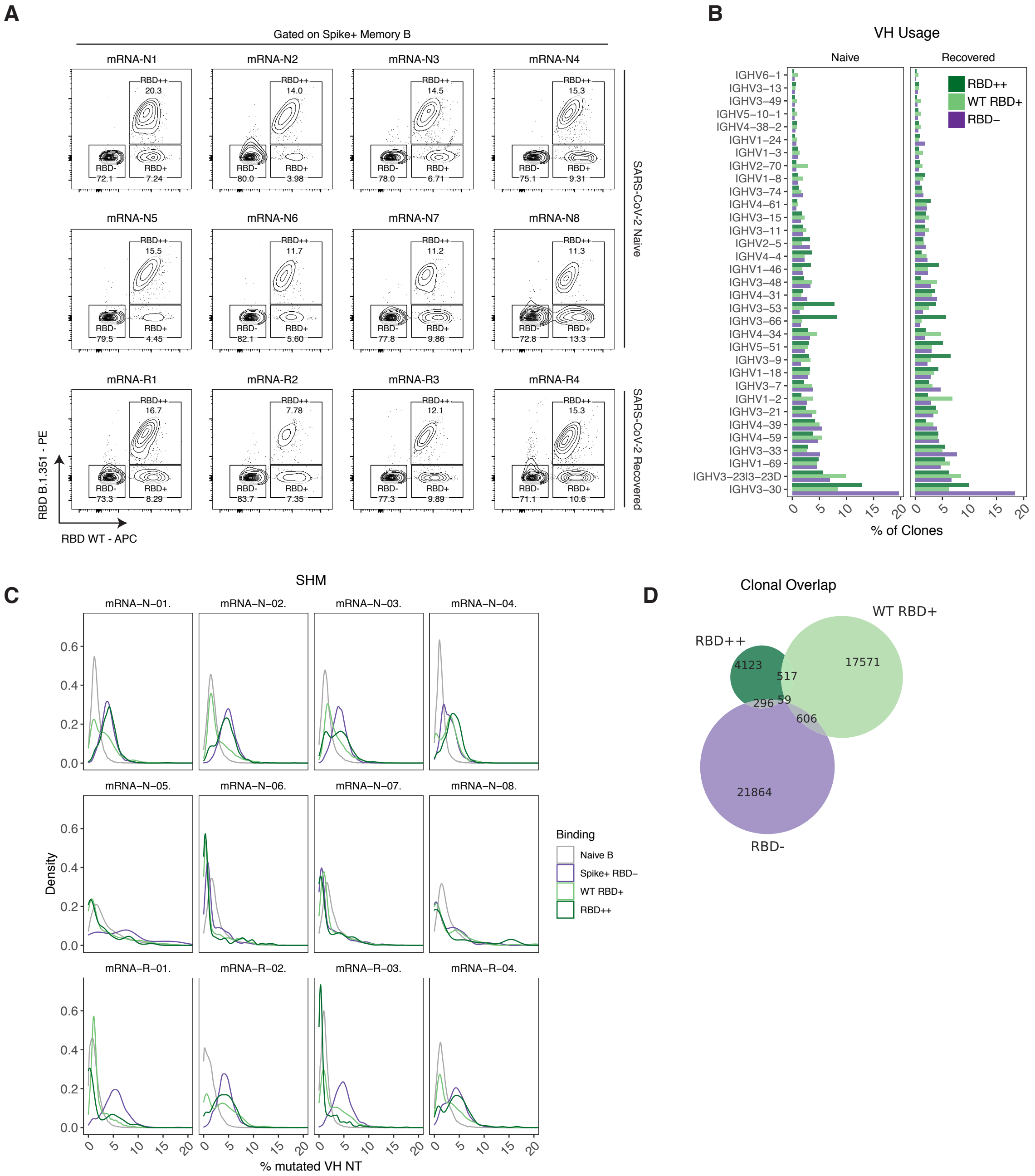
A
Gated on Spike+ Memory B
mRNA-N1
mRNA-N2
mRNA-N3
mRNA-N4
mRNA-N5
mRNA-N6
mRNA-N7
mRNA-N8
mRNA-R1
mRNA-R2
mRNA-R3
mRNA-R4
SARS-CoV-2 Naive
SARS-CoV-2 Recovered
RBD B.1.351 - PE
RBD WT - APC
B
VH Usage
Naive
Recovered
RBD++
WT RBD+
RBD−
% of Clones
C
SHM
Density
% mutated VH NT
Binding
Naive B
Spike+ RBD−
WT RBD+
RBD++
D
Clonal Overlap
RBD++
WT RBD+
RBD-
4123
517
17571
296
59
606
21864

### Figure S5

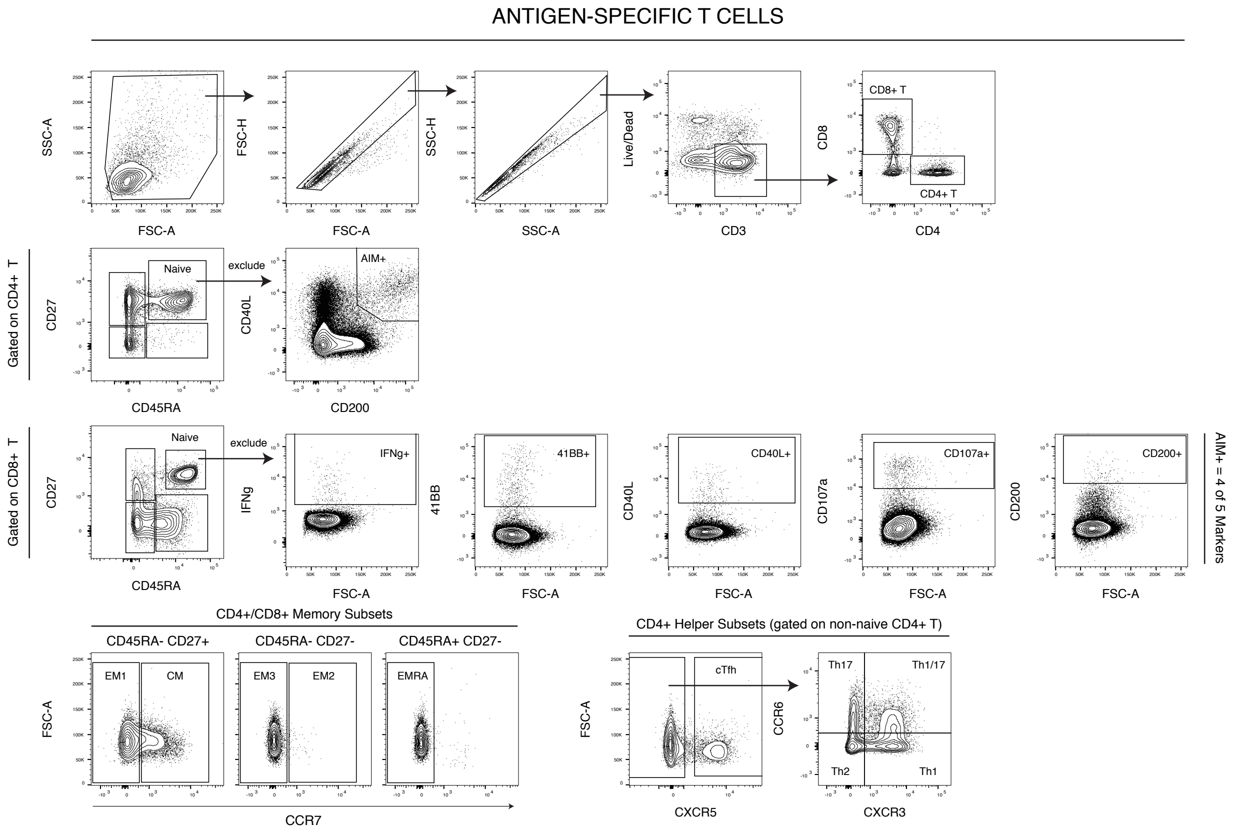

ANTIGEN-SPECIFIC T CELLS
SSC-A
FSC-A
FSC-H
SSC-H
Live/Dead
CD3
CD8
CD4
CD8+ T
CD4+ T
Gated on CD4+ T
CD27
CD45RA
Naive
exclude
CD40L
CD200
AIM+
Gated on CD8+ T
IFNg
IFNg+
41BB
41BB+
CD40L+
CD107a
CD107a+
CD200+
AIM+ = 4 of 5 Markers
CD4+/CD8+ Memory Subsets
CD45RA- CD27+
CD45RA- CD27-
CD45RA+ CD27-
EM1
CM
EM3
EM2
EMRA
CCR7
CD4+ Helper Subsets (gated on non-naive CD4+ T)
cTfh
CXCR5
CCR6
Th17
Th1/17
Th2
Th1
CXCR3

### Figure S6

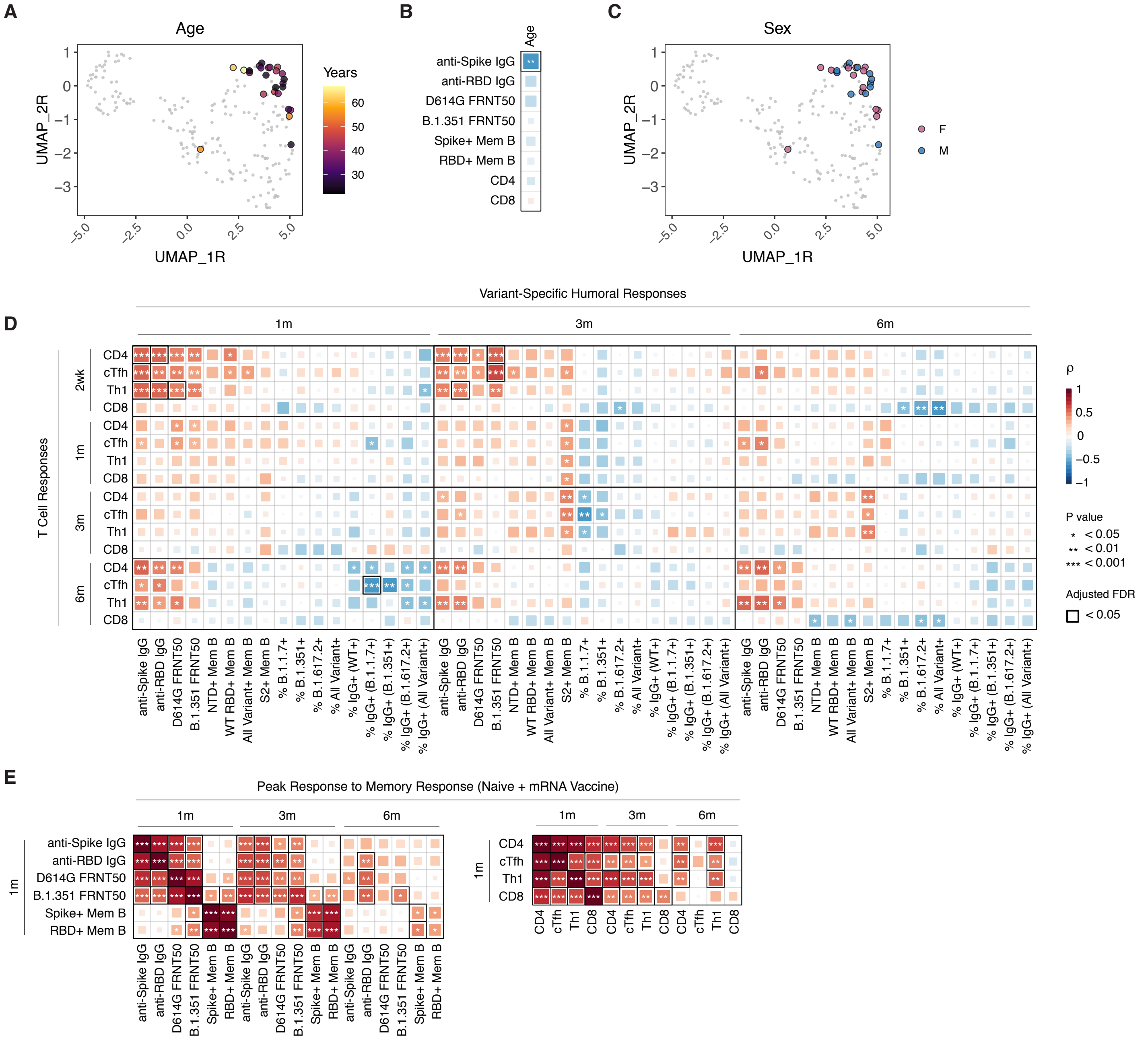
A
Age
Years
UMAP_2R
UMAP_1R
B
Age
anti-Spike IgG
anti-RBD IgG
D614G FRNT50
B.1.351 FRNT50
Spike+ Mem B
RBD+ Mem B
CD4
CD8
C
Sex
F
M
D
Variant-Specific Humoral Responses
1m
3m
6m
T Cell Responses
2wk
CD4
cTfh
Th1
CD8
anti-Spike IgG
anti-RBD IgG
D614G FRNT50
B.1.351 FRNT50
NTD+ Mem B
WT RBD+ Mem B
All Variant+ Mem B
S2+ Mem B
% B.1.1.7+
% B.1.351+
% B.1.617.2+
% All Variant+
% IgG+ (WT+)
% IgG+ (B.1.1.7+)
% IgG+ (B.1.351+)
% IgG+ (B.1.617.2+)
% IgG+ (All Variant+)
ρ
P value
Adjusted FDR
E
Peak Response to Memory Response (Naive + mRNA Vaccine)
Spike+ Mem B
RBD+ Mem B
